# The Neanderthal population history and the introgression landscape inferred from the UK Biobank

**DOI:** 10.64898/2026.04.03.716297

**Authors:** Adeline Morez Jacobs, Anahita Soltantouyeh, Roberta Zeloni, Francesca Carollo, Massimo Mezzavilla, Davide Marnetto, Luca Pagani

## Abstract

Neanderthal haplotypes in present-day Eurasians are unevenly distributed across the genome, forming introgression deserts and high-frequency segments consistent with adaptive introgression, with additional random variation affected by genetic drift. However, current estimates are limited by modest sample sizes and analyses restricted to subsets of the genome, given that any individual carries only 1-2% Neanderthal ancestry. Here we extract and analyse Neanderthal haplotypes from 45,000 imputed and phased genomes in the UK Biobank. Even at this scale, the number of sites overlapping Neanderthal haplotypes approaches—but does not reach—saturation, with rare haplotypes still being discovered. Using the derived allele frequency spectrum within the surviving Neanderthal segments, we infer a divergence time of 2,061 generations between the introgressed lineage and the Vindija Neanderthal, and estimate the effective population size of the introgressed lineage to Ne = 6,564. Individual-level resolution allows identification of 545 independent loci with excess Neanderthal homozygosity, consistent with ongoing selection. Despite the extensive dataset, a substantial portion of the genome remains a Neanderthal desert. Within these regions, we detect seven Human Accelerated Regions affected by recent human selective sweeps (TMRCA <650 kya), four located within introns of cerebellum-expressed genes, providing further support for their potential as modern human-specific adaptation.

## Introduction

Since the publication of the first draft of the Neanderthal genome (*1*), several admixture events between archaic and modern humans have been described (*2–4*). The most widespread event being the introgression from Neanderthal into all non-Africans as they expanded outside Africa, dated to around ∼50-45 kya (*5*, *6*), with more recent arrival in some African groups following back-to-Africa migrations (*7*). Today, non-African populations share ∼1-2% ancestry directly inherited from Neanderthals. The present-day proportion and distribution of Neanderthal ancestry is affected by antagonist evolutionary processes. Rapid purifying selection following introgression (*6*, *8*) purged 7-40% of the genome from Neanderthal variation (*9–12*), resulting in large regions significantly depleted of archaic ancestry―that can reach up to ∼10 Mb (*9–11*). This phenomenon was at least partially promoted by the accumulation of deleterious variants along the Neanderthal lineage caused by lower effective population size (Ne) (*13*). Moreover, sporadic cases of adaptive introgression conferred an advantage to the modern human populations expanding across Eurasia (*14–16*), and some of the causal variants reach frequencies 40-70% in some Eurasian groups (*16–18*). Nevertheless, a sizable portion of the Neanderthal contribution provided no effect to the modern human genome and, as such, remained invisible to natural selection. Past efforts to characterise the Neanderthal genome embedded within the genome of non-Africans have stressed its functional impact and contribution to the architecture of certain complex phenotypes (*19–21*). It has also been reported that European populations harbour more Neanderthal diversity than East Asian ones, reflecting the post-introgression population dynamics that characterised modern human groups (*22*).

Despite major advances, the data available to infer Neanderthal’s population history and impact of introgression remain limited to three high-coverage genomes (*23–25*), eight low-coverage nuclear genetic material (*26–28*). In addition, indirect reconstruction of surviving Neanderthal fragments within modern humans is mostly limited by analytical strategy that captured only a fraction of the Neanderthal variation segregating in present-day genomes (*20*, *29*).

We here leverage the availability of thousands of human imputed and phased genomes from the UK Biobank of European descent to i) reconstruct the full site frequency spectrum as well as landscape of private mutations of the introgressed Neanderthal genome, ii) refine the evolutionary history of the introgressed Neanderthals, iii) characterise patterns of ongoing selection and iv) refine our current understanding on the extent and pervasive nature of archaic deserts along the genome of modern Europeans.

## Results and Discussion

### Recovery of Neanderthal segments within 45,000 European genomes from the UK Biobank

We identified Neanderthal introgressed segments in 45,532 UK Biobank genomes, selected for having MRI and other brain-related phenotypes as described in (*30*). Introgressed segments were defined as regions showing a significant enrichment of derived alleles relative to Yoruba genomes, using the S* statistic implemented in SSTAR (*31*), followed by additional quality-control filtering (see Methods). Unlike reference-based approaches, SSTAR does not rely on a Neanderthal genome, providing greater sensitivity to archaic variation absent from currently sequenced Neanderthals. Our method recovered a total of 5,258,470 polymorphic positions covered by at least one Neanderthal segment, representing a ∼11.3% increase compared with 500 genomes on average—the sample size typically used in previous studies (Fig. 1a; (*9*, *10*, *32*). As the number of genomes increases, the rate of discovery of new variants within Neanderthal introgressed segments decreases but does not reach a plateau. This indicates that very rare variants associated with introgressed segments remain to be detected. Together, these results highlight the importance of large-scale genomic datasets for capturing rare Neanderthal-associated variation. We additionally tested whether some identified Neanderthal segments might still contain modern human sequences using SSTAR match rate and *f_4_-ratio*. SSTAR match rate calculates, for each segment, the proportion of variants marching either a Yoruba or the Altai Neanderthal genome. We also computed the *f_4_-ratio,* 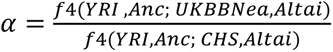 which estimates the proportion of gene flow (𝛼) from a modern human source (CHS, one Han Chinese genome) into the Neanderthal introgressed segments (UKBBNea). Conceptually, this ratio compares the excess genetic drift between a Yoruba genome and introgressed Neanderthal segments (numerator) to the full genetic drift between a Yoruba and a Han Chinese genome (denominator, see Methods). In this context, the *f_4_-ratio* can be interpreted as the fraction of falsely inferred introgressed segments. Overall, the inferred Neanderthal segments are robust to false discovery, with minimal residual modern human contributions, as shown in Fig. 1b (match rate) and c (*f_4_-ratios for each autosome*).

**Fig 1.**
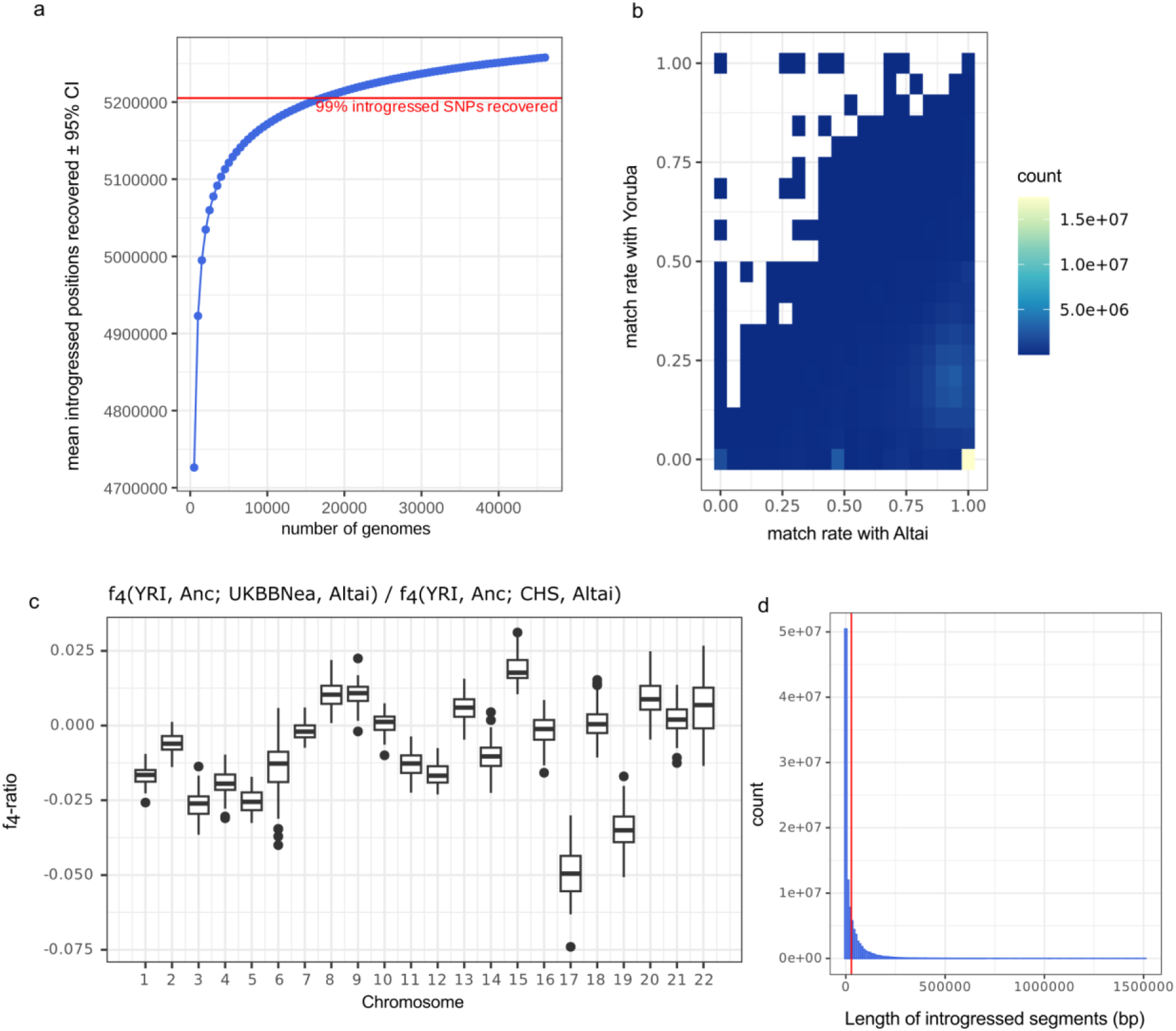
Neanderthal introgressed haplotypes in ∼45k UK Biobank genomes. **(a)** The recovery rate of new SNPs overlapping an introgressed haplotype decreases with sample size, but does not reach saturation (mean ± 95% CI over 100 iterations). On average, 99% of positions matching an introgressed haplotype are recovered with 17,500 genomes (red line). **(b)** The introgressed haplotype match rate is consistently higher in the high-coverage Altai Neanderthal genome than in the Yoruba population. **(c)** per-chromosome *f_4_-ratio* estimates across 92 batches of 1000 UKBB target haploid genomes. *f_4_-ratio represents* the proportion of admixture (α) in the Neanderthal introgressed segments from a modern human souce (represented here by a Han Chinese genome), which can be explain if false positive introgressed segments are still present. The highest *f_4_-ratio* for a batch α = 3.7% is visible in chromosome 15. YRI is a Yoruba genome which is used as the sister group of CHS, a Han Chinese genome. Anc is the ancestral genome, used as the outgroup in this phylogeny. **(d)** The length distribution of the introgressed Neanderthal segments, with an average of 28,082 bp (red line).

Using a working dataset restricted to 38,406 UK Biobank non-related samples of European ancestry (*30*), this approach yielded a collection of ∼102 million segments encompassing 5,113,265 polymorphic positions covered by at least one Neanderthal segment. We found an average Neanderthal ancestry proportion of 1.11% ± 0.09% standard deviation (SD) per individual, consistent with previous estimates from alternative methods (*9*, *32*). Importantly, the reported ancestry proportion is based solely on identifiable archaic segments, which altogether make up for a smaller fraction of the total Neanderthal contribution to the genomes of non Africans.

### The evolutionary history of the introgressed Neanderthal lineage

Thanks to this extensive dataset, we can characterise the site frequency spectrum (SFS) of variants carried within Neanderthal haplotypes inherited by present-day humans (Fig. 2a). We supplied this SFS as input to *fastsimcoal* 2.8 (*33*, *34*) to infer demographic processes that affected specifically the introgressed Neanderthal lineage―otherwise unsampled―, before and shortly after the introgression event. We were particularly interested in its divergence time from the three high-coverage Neanderthals and in its effective population size. The SFS was restricted to sites considered unaffected by background selection and biased gene conversion, following recommendations in (*35*). Because the introgressed Neanderthal SFS is also shaped by demographic events affecting the modern human lineage after Neanderthal introgression, we developed an innovative approach to input the demographic history of the European UKBB population (Fig. 2b) into the UKBB Neanderthal simulations (Fig. 2c). Demographic modelling of the European UKBB, Yoruba, and the three high-coverage Neanderthals provided good fits to the data (Figs. S1-S3). Replacing the European UKBB with the Tuscan genomes (TSI) from the 1000 Genome Project produced nearly identical inferences, indicating that our results are robust to differences in data generation (Fig. S4).

**Fig 2.**
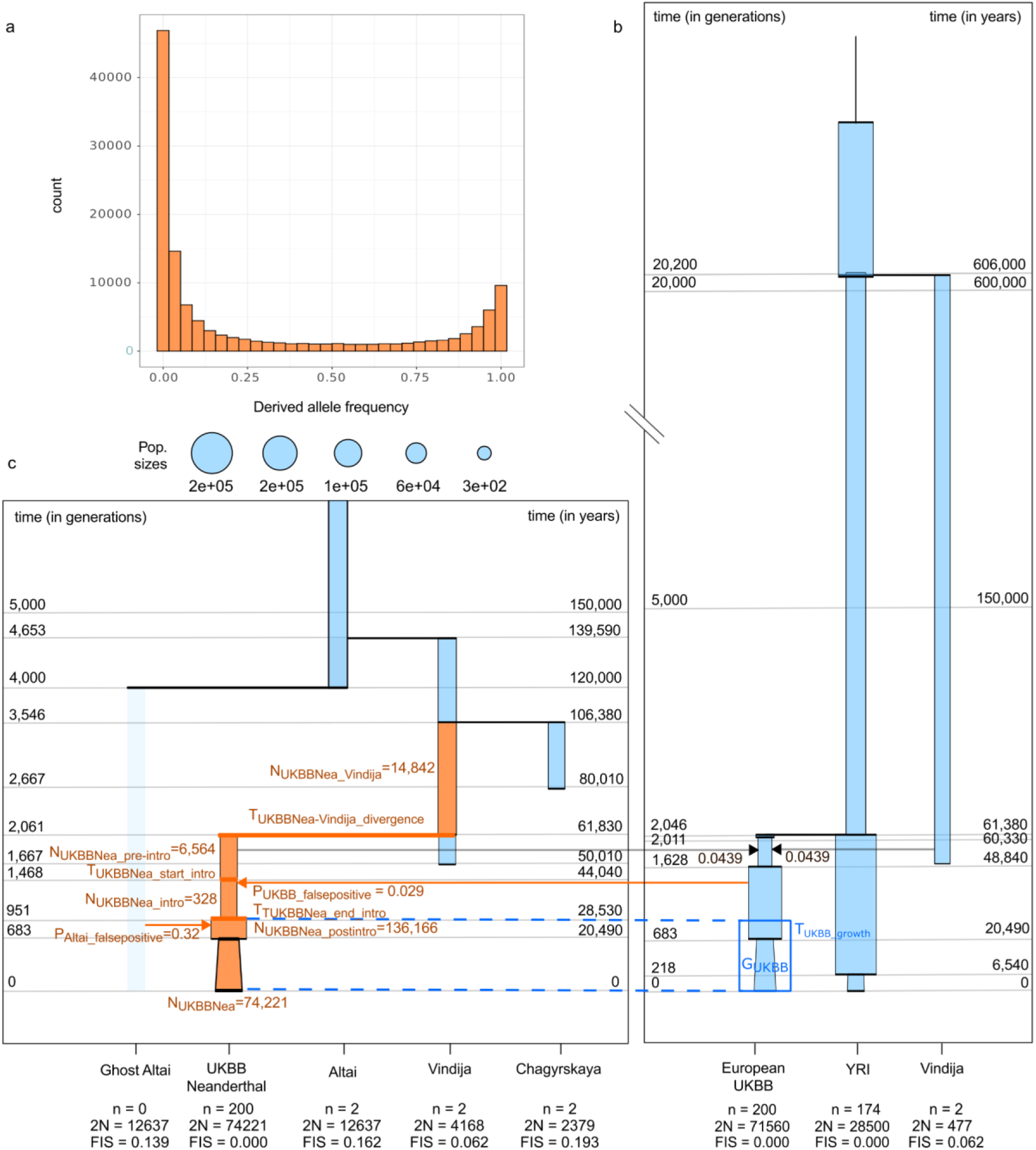
Demography modelling of the introgressed Neanderthal lineage. The derived allele frequency (DAF) spectrum within the introgressed Neanderthal segments **(a)** was used to infer the demographic history specific to the introgressed Neanderthal lineage **(c)**. The inference explicitly incorporates the post-introgression demographic history of European UKBB populations, inferred as in **(b)** to account for demographic processes affecting the observed DAF spectrum after introgression (Table S1). Specifically, modern UKBB’s growth rate (*G_UKBB_*) and timing (*T_UKBB_growth_*) was used (in dark blue), while the exact population size was freely simulated by *fastsimcoal* as it will still be impacted by Neanderthal’s past demography. Inferred parameters in (c) are highlighted in orange (Table S5). The starting time of the modern UKBB demography depends on the inferred end of the introgression bottleneck, which was freely simulated by *fastsimcoal* (*T_UKBBNea_end_intro_*; Table S5). Gene flow from the Altai Neanderthal and the European UKBB into the introgressed Neanderthal lineage was explicitly added into the simulation to account for false positives in the detection of introgressed segments and the use of the Altai genome in the ascertainment of the introgressed haplotypes (orange arrows). The migration rates were fitted by the simulations. See Figs. S1 and S5 for the fit between observed and expected DAF spectrum. n represents the sample size, 2N represents the effective population size, and FIS represents the inbreeding coefficient.

We next incorporated the introgressed Neanderthal lineage into a joint model of archaic and modern humans. We used the estimated European UKBB parameters (Fig. 2b)ーespecially the growth rate *G_UKBB_* and the onset time of growth *T_UKBB_growth_* (Table S1)ーto model the post-admixture evolution of the introgressed Neanderthal segments (Fig. 2c). However, we allowed the exact effective population sizes pre- and post-growth (N_UKBBNea_postintro_ and N_UKBBNea_, Table S5) to be freely inferred in *fastsimcoal,* because Neanderthal demographic history may influence these parameters differently than in the European UKBB population. Allele-based trees using *TREEMIX* (*36*) and *qpGraph* (*37, 38*) confirmed a higher affinity of the introgressed lineage to Vindija than to Altai or Chagyrskaya (Figs S6 and S7; (*24*). We accounted for potential biases in our introgression detection methodology—including over-assignment of fragments matching Altai-specific variants and a low rate of false-positive introgressed segments that are actually modern humans—by allowing migrations from both Altai and UKBB into the introgressed lineage; this choice is also supported by allele-based trees (Figs. S6 and S7).

Our best-fitting *fastsimcoal* model estimates a divergence time between the introgressed Neanderthal lineage and Vindija of T_UKBB-Vindija_divergence_ = 2,061 ± 160 generations (Observed Maximum Likelihood (ML) ±95% block-bootstrapped confidence interval (CI)), corresponding to approximately 61.8 ± 4.8 kya, assuming a 30-year generation time (Observed ML = - 46,466,555.244, Δ_expected ML-observed ML_ = 165,888, Fig. 2a). This date closely pre-dates the timing of Neanderthal introgression into modern humans in our model T_intro_ = 55.6 ± 3.3 kya (ML ±95% CI, Fig. 2b, Table S1) and with previous estimates of 50.5-43.5 kya (*6*), confirming that Vindija is a near-direct representative of the Neanderthal population that admixed with early non-Africans (*24*). The inferred divergence also provides an upper bound for the split between the introgressed lineage and the low-coverage Neanderthal from Mezmaiskaya in the Caucasus, which is estimated to be even more closely related to the admixing population (*23*). Together, these results are consistent with the hypothesised southwestern Asia region for Neanderthal-modern human interbreeding, at the southernmost extent of the late Neanderthal range. Integrating genetic and palaeoecological data, a recent study proposed the Persian plateau as a particularly plausible region for contact, as it may have acted as a long-term demographic hub for modern humans for up to ∼30 ky, between the Out of Africa (∼70-60 kya) and the later dispersal of which all present-day Europeans descend (∼38 kya, (*39*). Archaeological evidence from Bawa Yawan (*40*) and Wezmeh (*41*) further indicates that Neanderthals occupied this region at the time of the modern human Out of Africa. Additional episodes of introgression likely occurred in Europe, with the early modern humans from Oase (42-37 kya) and Bacho Kiro (45.9-42.6 kya) long Neanderthal tracts reflecting admixture within the previous 4–17 generations (*42*, *43*). However, the uniqueness of their Neanderthal haplotype, together with the subsequent replacement of these early populations, implies that these late European contacts made little or no contribution to the Neanderthal ancestry found in present-day Europeans (*6*).

The effective population size of the introgressed Neanderthal lineage after its split from Vindija is inferred to be small (N_UKBBNea_pre-intro_ = 6,564 ± 1,100, ML ±95% CI), falling on the upper limit of the range estimated for Vindija (N_Vindija_ = 4,168 ± 1,580), and Chagyrskaya (N_Chagyrskaya_ = 2,379 ± 1,845) (*24*, *25*, *44*) Our estimate for Altai (N_Altai_ = 12,637 ± 2,265) is slightly higher than previous reports (*23*, *24*, *44*). Our method, which explicitly considers reduction of diversity in the introgressed Neanderthal due to admixture bottleneck and subsequent purifying selection, yielded a slightly higher N_UKBBNea_pre-intro_ estimate than in (*21*). Interestingly, the Ne estimated for the introgressing Neanderthal population is slightly higher than our estimate for the European population Ne during the Out of Africa (N_OoA_ = 3,124 ± 813), comparable with previous estimates (*45*, *46*), suggesting that the Neanderthal populations that encountered the modern humans were not so small in comparison. This resonates with findings pointing to a higher inter-band connectivity later on at ∼38 kya to explain the successful dispersal of humans (*47*), instead of being caused by the immediate absolute population density. Finally, we infer a prolonged bottleneck in the introgressed Neanderthal lineage, spanning the introgression event and the subsequent purging of deleterious variants, from T_UKBBNea_start_intro_ = 44.0 ± 4.2 kya to T_UKBBNea_end_intro_ = 28.5 ± 2.6 kya, and involved at least N_UKBBNea_intro_ = 328 ± 55 Neanderthal individuals, although this estimate is likely downward biased by both above-mentioned mechanisms.

### Expanding the set of Neanderthal-specific variants

To characterise the functional consequences of adaptive introgression, we first refined the set of variants likely to be of genuine Neanderthal origin, excluding alleles shared with the common ancestors of both modern humans and Neanderthals, as well as mutations arising on the modern human lineage after introgression. As a high-confidence reference set, we defined Neanderthal-specific variants as those observed in at least one of the three high-coverage Neanderthal genomes and at <1% frequency in the Yoruba population (n = 346,401 variants, Fig. 3a, in blue).

We then implemented a novel inference framework to identify additional Neanderthal-derived variants that are absent from currently sequenced archaic genomes. Because genuinely introgressed alleles are expected to remain in strong linkage with the Neanderthal-introgressed haplotypes, their DAF on these segments should exceed that of post-introgression mutations. We therefore used the empirical DAF distribution of observed Neanderthal-specific alleles on introgressed haplotypes to infer additional Neanderthal-private candidate variants absent from the three high-coverage Neanderthal genomes, and with a DAF <1% in the Yoruba. Retaining variants within the upper 95% of this distribution (DAF > 0.69, Fig. 3a), we identified 50,680 additional derived variants strongly associated with introgressed segments. This strategy substantially expands the catalogue of likely Neanderthal-derived variation beyond the limited number of sequenced archaic genomes. We also note that the logic of our approach is akin to the LD-based one described in (*29*)but in our case we constrained the additional Neanderthal variants to fall within an introgressed haplotype.

### Evidence for ongoing selection of introgressed variants

We next asked whether any of these introgressed alleles exhibit signatures of ongoing or recent positive selection. Leveraging individual-level archaic load in the UK Biobank, we implemented a Hardy–Weinberg equilibrium (HWE)-based framework to detect genomic regions exhibiting an excess of introgression homozygotes (see Methods). Unlike traditional selection scans that rely on long-term allele frequency shifts, HWE-based approaches are particularly sensitive to ongoing selection because they directly capture distortions in genotype proportions (*48*). This framework therefore enables detection of both persistent positive selection acting since the admixture and more recent adaptive responses to changing environments, including weak positive selection acting on alleles at low-to-moderate frequency.

We identified 2,076 positions displaying significant Hardy-Weinberg disequilibrium (p < 10^-8^, Table S6) characterised by an excess of introgression homozygotes, carrying at least one Neanderthal-specific derived allele. After clumping (r^2^ > 0.2 within 100 kb), 567 independent signals remained (Fig. 3b). In order to retain true candidates of adaptive introgression, we excluded variants whose HWE deviation was likely driven by a non-introgressed position in LD (r^2^ > 0.2, ± 100 kb from the introgressed variant), showing a stronger association (i.e. lower p-value) than the introgressed variant. Following this filtering step, 545 positions remained as putative target of positive selection.

Strikingly, the signals span a broad allele-frequency spectrum, including variants with DAF < 10%. Overall, a majority of variants (56.3%) show strong European-African differentiation, falling within the top 90^th^ percentile of F_ST_ between GBR and YRI, consistent with adaptive introgression, and only 43 variants overlap with signal of adaptive introgression from Neanderthal previously reported in (*49*). This highlight the power of our approach to detect subtle but ongoing selective pressures. For 348 positions, the Neanderthal-specific derived allele is the only introgressed allele, while 197 show residual-to-high variability (Fig. 3b, orange shades; Table S6). We observed a residual-to-moderate but widespread underestimation of the introgression rate of Neanderthal-specific derived allele in the UKBBEUR compared to its DAF in the UKBBEUR (10.8 ± 9.3% (mean difference ± SD); Table S6). This discrepancy suggests that some introgressed segments might not be detected, but we cannot exclude that, in some instances, the derived allele was incorrectly attributed as Neanderthals-specific. The latter could occur if the allele was recently lost in the Yoruba lineage and actually represents a variant inherited from the common ancestor shared by modern humans and Neanderthals. Among the 545 independent variants, 21 are not observed in any high-coverage Neanderthal (red dots in Fig. 3b). Three of these exhibit CADD PHRED scores > 10, suggesting functional relevance (rs71446761, PHRED = 14.27; rs1316763, PHRED = 13.13; rs74477231, PHRED = 12.72; Table S6). Notably, rs71446761 is located within *SLCO1B1*, a liver-expressed gene (*50*). While these variants can reflect long-standing polymorphism within Neanderthal that are unsampled in the high-coverage genomes, they can also underline the functional impact specific to the introgressed Neanderthal lineage.

Variants in significant HWE deviation and with high DAF (>0.7, Fig. 3b) overlap coding regions of *OR9G1* and *PPDPF*, previously implicated in adaptive introgression (*49*, *51*), in the exon of *STARD9,* for which the underlying evolutionary process is discussed (*52*). Additional signals are localised in the intron of *NECAB1*, and upstream *VEPH1* and *RSRC1*, expanding the landscape of candidate loci potentially shaped by Neanderthal ancestry, to our knowledge.

To infer the functional implications of the variants in significant HWE deviation, given that MAF and LD-score have a significant effect on the power to retrieve significant genotype-phenotype associations (*53–55*), we first created a null genomic background matching DAF and LD-score (defined as r^2^ sum over 200-kb window) distribution as a reference set (see Methods). We then tested for enrichment in hits resulting from Genome Wide Association Studies (GWAS) and expression Quantitative Trait Loci (eQTLs) against this null background. The 545 independent signals overlap 33 unique GWAS associations catalogued in the NHGRI-EBI GWAS database ((*56*); Table S7), although this overlap does not exceed expectation relative to the null genomic background (p = 0.216). Likewise, when looking for an enrichment in tissue-specific eQTLs from GTEx (*57*), no tissue reached significance after multiple testing correction (threshold = 1.02x10^-3^). However, the top nominal enrichment was observed in cultured fibroblasts (p = 0.019; Table S8).

The 545 LD-clumped variants enriched in Neanderthal introgressed homozygotes show significant enrichment across eight Gene Ontology (GO) biological categories and five mouse phenotype categories, with nine categories represented by at least five independent regions (FDR-corrected q-value < 0.05 and fold enrichment ≥ 1.5; Fig. 3c), relative to a DAF- and LD-score-matched genomic background. These enrichments converge on five broad biological domains: immune regulation (negative regulation of B cell activation), cellular stress-response activity (response to cobalt ion), intracellular transport (Golgi to lysosome transport, post-Golgi vesicle-mediated transport), craniofacial and musculoskeletal morphology (absent hard palate, abnormal pectoral muscle morphology) and neural/behavioural phenotypes (glial cell apoptotic process, social withdrawal). This convergence aligns with prior findings that Neanderthal introgression contributed to facial morphology, by increasing nasal and midface height (*58*) and modulating expression near genes impacting the cartilage (*59*), the adaptive immune system (*60*, *61*), the nervous system, including specifically variants near *PHLPP1* involved in glial apoptosis (*62–64*). Our results extend these observations by demonstrating that a subset of these introgressed alleles continues to exhibit genotype-level distortions consistent with ongoing selection in contemporary populations. Together, these findings suggest that the adaptive legacy of Neanderthal admixture remains evolutionarily dynamic, with some introgressed alleles maintained, and potentially favoured, in the present-day environment.

**Fig 3.**
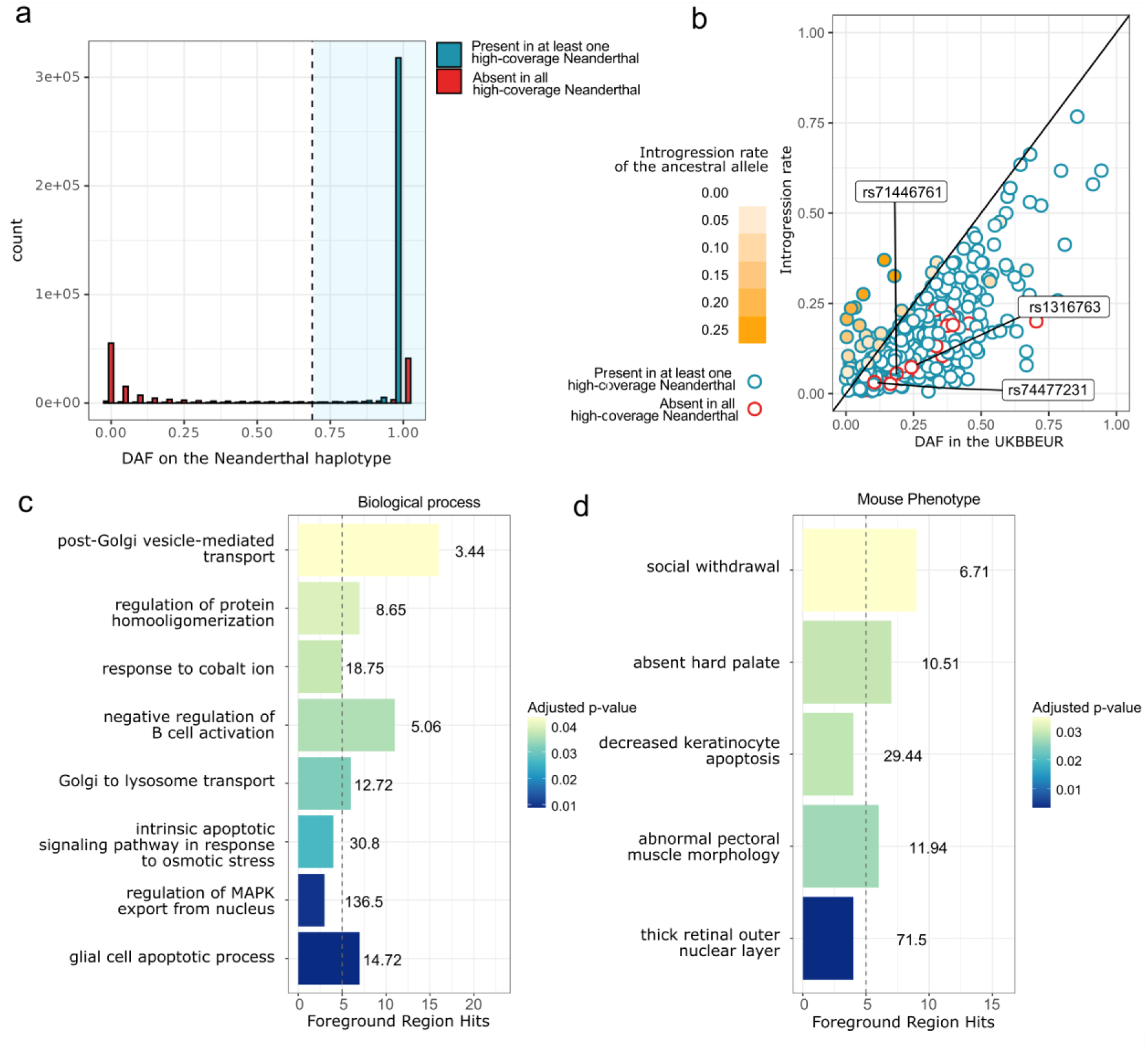
Neanderthal-introgressed variants in deviation from Hardy-Weinberg equilibrium (HWE) and their functional impact in the European UKBB. **(a)** Derived Allele Frequency (DAF) distribution of observed Neanderthal-specific variants (present in at least one high-coverage Neanderthal genome and <1% in the Yoruba, in blue) and DAF distribution of variants absent in all high-coverage Neanderthal genomes and <1% Yoruba (in red). We used the upper 95% of the distribution of Neanderthal-specific variants (within the blue-shaded box) to infer putative Neanderthal-specific variants not observed in the high-coverage archaic genomes (DAF > 0.69). **(b)** Introgression rate and DAF in the European UKBB of observed (blue dots) and putative (red dots) Neanderthal-specific variants with a significant deviation from HWE for an excess of Neanderthal-introgressed homozygotes, after clumping (r^2^ > 0.2 within 100 kb). The orange gradient indicates the introgression rate of the ancestral allele frequency. The darker the colour, the higher the frequency of the ancestral allele in the introgressed segments, indicative of Neanderthal variability. Deviation between introgression rate and DAF in the UKBBEUR can be explained either by a lower introgression detection power, or inheritance from the modern human-Neanderthal common ancestor, recently lost in the Yoruba lineage. Putative Neanderthal-specific variants with a functional relevance (CADD PHRED score > 10) are annotated. **(c)** and **(d)** Enriched GO categories after correction (FDR-corrected p-value < 0.05) of the Neanderthal-specific variants in (b). Numbers on the bars indicate the fold enrichment, and bar height represents the number of foreground region hits. The adjusted p-values are represented by the colour scale, with darker shades indicating smaller (more significant) p-values. The grey dashed vertical line indicates the foreground region hit threshold ≥ 5.

### A refined map of the Neanderthal desert

We defined regions of the genome depleted of Neanderthal introgression as 10-kb windows in which the mean inferred Neanderthal introgression rate is <10⁻³ and in which no site exhibited an introgression rate ≥10⁻². These windows were further classified into two categories: (i) windows containing at least one Neanderthal-specific SNP (strict gaps); and (ii) windows containing no Neanderthal-specific SNPs (ILS), which likely reflect Incomplete Lineage Sorting (ILS), with a limited power to discriminate between genuine introgression and lack thereof. Strict gaps comprise 13.8% of the European UKBB genome (Fig. 4a, light), comparable to a previous ARG-based estimate, explicitly considering ILS (*12*). Given that, on average, only ∼2.4 mutations per 10-kb segment are expected to have accumulated on the Altai Neanderthal lineage since its split with modern humans (see Methods), many 10-kb windows are expected to lack Neanderthal-specific mutations. Consequently, a substantial fraction of true Neanderthal depleted windows are classified as ILS by this criterion. To obtain a more permissive definition of introgression deserts, we assigned ILS windows to either gaps or introgressed when they were flanked on both sides by the same category, and refer to them and their strict counterpart as expanded gaps and expanded introgressed regions, respectively. Despite the size of the present dataset, 35.1% of the genome remains devoid of Neanderthal introgression (Fig. 4a, dark), indicating that previously reported introgression deserts cannot be attributed to limited sample size (*9*, *12*, *32*).

**Fig 4.**
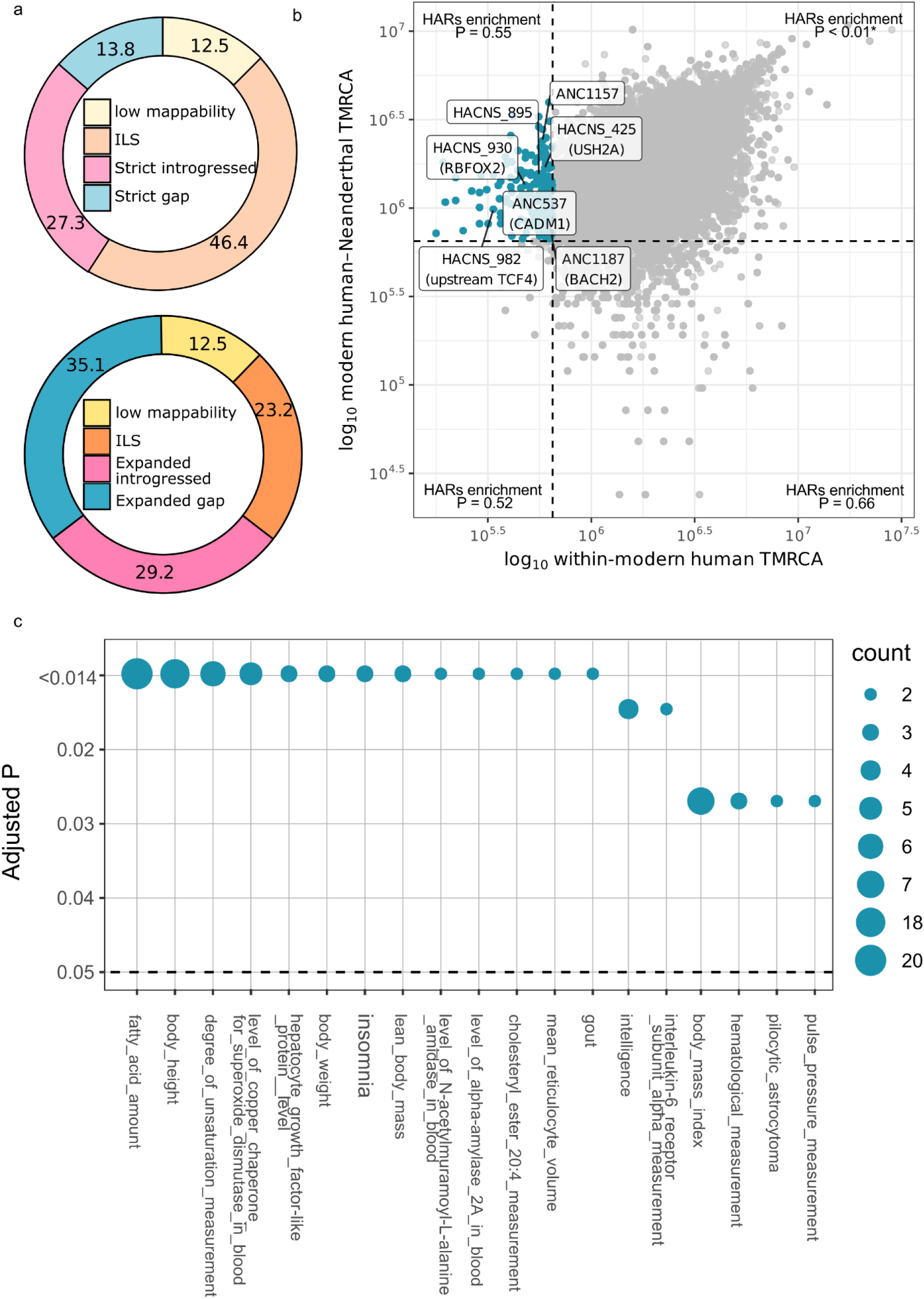
Description of the Neanderthal introgression depletion. **(a)** Genomic proportions of introgressed regions and regions depleted of Neanderthal introgression, shown before (light), and after (dark) assigning ILS windows flanked on both sides to gap or introgressed. **(b)** expanded desert partitioned by their TMRCA within modern humans and between modern humans and Neanderthals. Dashed lines represent 650 kya, the higher estimate for the split between Neanderthals and modern humans. Windows associated with a modern human-Denisova TMRCA <650 kya were excluded to control for the region under long-lasting purifying selection. We explored the enrichment for the Human Accelerated Regions (HARs) into each category, and Benjamini-Yekutieli FDR-adjusted P-values were reported. We focused on the expanded gaps with a recent TMRCA within modern humans (<650 kya) but with a modern human-Neanderthal TMRCA ≥ 650 kya (blue dots), indicative of a recent sweep specific to the modern human lineage. The seven matching HARs are annotated, including the associated genes. **(c)** GWAS catalogue EFO traits significantly enriched in introgression gaps with a recent modern human TMRCA, at Benjamini-Yekutieli FDR-adjusted P-value < 0.05 and with number of GWAS catalogue hit (count) > 1.

We show that the expanded gaps are, on average, more evolutionarily constrained and intolerant to genetic variation than the expanded introgressed windows or the remaining ILS windows (Table S9), extending earlier observations that Neanderthal introgression is preferentially excluded from functionally important parts of the genome (*9*). Expanded gaps are also more frequently involved in protein-protein interactions than introgressed windows. Expanded gaps occur in regions of significantly lower recombination rate than the expanded introgression (Wilcoxon test, p < 10^-15^), indicating that the depletion of Neanderthal ancestry is not an artefact of reduced power to detect introgression in regions of high recombination. Instead, it might reflect the tendency of introgression gaps to be in specific genomic regions associated with lower recombination rate (*65*, *66*).

Three of the four previously reported long introgression deserts (*10*) fall within the top 5% of the 10-Mb windows enriched for expanded gaps, and two partially overlap the top 1% (chr3:77,000,000-89,000,000 and chr7:109,000,000-125,000,000; Table S10). We further identify long, uninterrupted expanded gaps across the genome, with the top 0.1% longest gaps comprising nine regions overlapping 98 genes (Table S11). Notably, these genes are enriched for the filopodium-associated GO term (GO:0030175, FDR-adjusted p = 0.0215). This enrichment is driven by six genes distributed across four distinct genomic regions, supporting a trans-acting biological signal rather than an artefact of local gene clustering (Fig. S11, Table S11).

### The evolutionary history of deserts

We next exploited the information contained within introgression gaps to construct a refined map of genomic regions that are selectively depleted of Neanderthal variation and specific to modern humans. To this end, we classified expanded gaps according to their time to most recent common ancestor (TMRCA) estimated within modern humans and between modern humans and Altai Neanderthal (*67*), partitioning them using a threshold of 650 kya corresponding to the modern humans-Neanderthal split (*23*) (Fig. 4b). If the within modern human TMRCA is > 650 kya it means that the presence of multiple haplotypes surviving in present-day humans predates this split, otherwise the present-day haplotype variability originates after the split. Assuming a long term Ne of 10,000 and a generation time of 30 years, the within modern human TMRCA is expected to be 4 Ne, or 1,200,000 years (64, 65). A significant decrease in the empirical within modern human TMRCA can be interpreted as evidence for purifying or positive selection, as well as caused by random effects such as genetic drift. To specifically exploit regions where Neanderthal variation has been purified and exclude highly conserved regions in the hominin lineage, we further excluded from the analysis windows with modern human-Denisova TMRCA < 650 kya. Strikingly, most expanded gaps fall into deeply divergent windows both within modern humans and between modern humans and Neanderthal (TMRCA ≥ 650 kya). This pattern indicates that introgression gaps are preferentially located in haplotypes that predate the divergence of the two lineages and is consistent with purifying selection acting to remove deleterious alleles likely accumulated along the Neanderthal lineage because of their reduced effective population size.

We further tested whether any genome category (expanded gap, expanded introgressed, ILS) is enriched for Human Accelerated Region (HARs) (*68*) (i.e. genomic elements that are highly conserved across vertebrates but show unusually rapid evolution on the human lineage). Although HARs must have accumulated after the human-chimpanzee divergence (∼6-8 Mya), their temporal distribution along the human lineage remains largely unresolved. We found that HARs are significantly enriched within the expanded gaps (permutation P < 0.001), but not in ILS (permutation P = 0.723) or introgressed regions (permutation P = 0.119). Importantly, when stratifying expanded gaps by TMRCA, this enrichment is entirely driven by the subset of gaps having both within-modern human and modern human-Neanderthal TMRCA ≥ 650 kya (permutation P < 0.001). If HARs had arisen more frequently than expected on the modern human-specific lineage, one would instead expect enrichment in gaps with shallow within-human TMRCA (< 650 kya), which we do not observe. These results indicate instead that a higher than expected fraction of HARs originated prior to the divergence of modern humans and Neanderthals, that variations still existed at least within Neanderthals, and that Neanderthal alleles at these loci were subsequently purged by purifying selection following admixture. However, we cannot exclude that higher observed TMRCA can result from standing variation persistence or recombination in the modern human lineage following the appearance of the functionally relevant mutation(s), resulting in a TMRCA predating the selection event. As support for an enrichment of HARs prior 650 kya, an Ancestral Recombination Graph approach, considering a more refined recombination map, also reported the observed pattern (*68*).

### The deserts with modern human-specific variation

We then focus on expanded gaps specific to modern humans (0.3% of the genome, Fig. 4b), which are defined as regions with a recent TMRCA < 650 kya but whose Neanderthal-modern human divergence predates the populations’ divergence (TMRCA ≥ 650 kya). These regions are particularly interesting, as they hold the power to detect modern humans genetic specificity that have been actively purified of Neanderthal introgression. We could replicate previous findings, showing that the expanded gaps with a recent modern human TMRCA includes parts of the *SPAG5* gene, involved in the mitotic spindle, and known to harbour 5 non-synonymous substitutions between modern humans and Neanderthals (*23*, *52*).

Next, we identified an overlap with seven HARs. Four of these fall within introns of *RBFOX2*, *CADM1*, *BACH2* and *USH2A*, and one is ∼27 kb upstream of *TCF4* (Table S12). Notably, *RBFOX2, CADM1,* and *TCF4* show their highest median expression in the cerebellum (*50*). They primary function is brain-related: *RBFOX2* encodes an RNA-binding protein that is thought to be a key regulator of alternative exon splicing in the nervous system and other cell types (*69*), *CADM1* (also known as SynCAM) is a cell adhesion molecule involved in the synapse formation and maintenance, and regulation of neurotransmitter release in the synapse (*70*) and *TCF4* encodes a transcription factor broadly expressed, and may play an important role in nervous system development, as mutations are associated with the Pitt-Hopkins syndrome, or different neurodevelopmental conditions such as autism and schizophrenia (*71–73*). *BACH2* is a transcriptional regulator expressed in several cell types. However, it is over-expressed in the human pre-birth brain (∼2.4 times more) and cerebellum (∼4 times more) relative to the macaque (*74*).

We next asked whether gaps that experienced a recent sweep in the human lineage are enriched for particular GWAS traits, under the assumption that residual variation within these gaps may reflect the phenotype target of selection. We found 19 traits significantly enriched in these swept gaps, including traits associated with cognition, body size measurements, and lipid metabolism (Fig. 4c; Benjamini-Yekutieli FDR-adjusted P < 0.05). However, we caution that this interpretation is limited, because the causal variant under selection may be fixed in modern humans, while the tagging polymorphic variants used in GWAS could influence phenotypes unrelated to the true target of selection. We did not detect any significant enrichment in expression regulation in any particular tissue (Table S13).

Overall, previous works on the functional impact of the Neanderthal desert pointed to an enrichment of expression in the brain, as well as in testis (*9–12*). In our study, we found three genes overlapping with HARs, having a recent modern human TMRCA and falling in a Neanderthal gap, showing an increased expression particularly in the cerebellum. Interestingly, this signal is consistent with the distinct set of genes reported in (*12*). The cerebellum plays an important role in functions decisive in human evolution and survival, as it is involved in motor control and balance, cognitive functions such as attention and language, and emotional control, including the regulation of fear and pleasure responses (*75–78*). This finding resonates with the now well-established observed difference between the endocast of modern human and Neanderthal. The posterior cranial fossa, housing the cerebellum, is rounded and expanded in modern human, and is responsible for the globularity of the modern human skull, along with bulging parietal lobe and a steep frontal (*62*, *79–81*). Modern human-specific globularity occurred gradually in the modern human lineage, to reach modern variation between 100-35 kya (*82*).

## Conclusion

Our analysis of Neanderthal introgressed segments across an extensive dataset of ∼45,000 UKBB genomes reveals an unexpected persistence of very low-frequency introgressed variants. By characterising the remaining diversity within these fragments through the derived allele frequency spectrum, and by explicitly modelling the post-introgression demographic history, we refined demographic parameters specific to the introgressed Neanderthal lineage. Our estimate of its effective population size is relatively small and consistent with those inferred from available high-coverage Neanderthal genomes and confirms the close genetic affinity between the introgressed lineage and the Vindija (Croatia) late Neanderthal population (divergence dated to ∼61.8 kya). Interestingly, the inferred effective population size of the introgressing Neanderthal lineage is comparable to that estimated for the human populations soon after they left Africa. This observation suggests that the demographic decline of Neanderthals relative to expanding modern human populations may not simply reflect overwhelming demographic replacement, but support a more complex scenario where cooler and dryer MIS3 can explain Neanderthal habitat fragmentation (*83*), weakening inter-band connectivity among Neanderthal groups, which ultimately led to their extinction, even when modern human fitness, population size and growth rate remain similar to that of the Neanderthals (*47*).

The unprecedented sample size of this study also allows us to refine current knowledge on Neanderthal deserts in the modern human genome, showing that they are not artefacts of limited sampling, and revealing the full extent of long and short deserts interspersed across the non African genome. Overall, our analysis highlights the two main selective forces acting on introgressed variation after admixture. Purifying selection against introgressed variants, which is quintessentially represented by deserts, is most prominent in regions associated with cognition, body size, and lipid metabolism. Conversely, regions showing adaptive retention of Neanderthal variants consistent with ongoing positive selective effect (identified through excess of introgressed homozygotes) are enriched for traits related to immune regulation, cellular stress response, craniofacial and musculoskeletal morphology, and neural or behavioural phenotypes. Although these domains are partially distinct, neural and behavioural traits appear in both categories, suggesting a complex and possibly bidirectional selective regime affecting brain-related phenotypes, as also suggested by the study of brain morphology (*30*).

By leveraging the coalescence time across genomic regions, we disentangle deserts created by purifying selection against variants that accumulated under reduced efficacy of selection in Neanderthals—likely due to their small effective population size— from a smaller subset that appear to have been recently acquired and strongly selected in the modern human lineage. Among these, we identify four genes (*RBFOX2, CADM1, TCF4, BACH2*) not previously highlighted in this context that show strong depletion of Neanderthal variation, are associated with HARs, and display enriched expression in the cerebellum. This is particularly intriguing, given the growing evidence linking cerebellar development to cognitive functions that are highly elaborated in modern humans (*75–78*).

Although the study of Neanderthal-modern human introgression can provide valuable insights into the evolutionary drivers underlying human specificity, the complex nature of these traits make establishing direct links between specific archaic/modern human variants and present-day phenotypes challenging (*29*, *84*). Overall, our results show the power of leveraging large population datasets to recover most of the available Neanderthal signature leftover in non African genomes, and pave the way to functional genomics and experimental perturbation study to better resolve the phenotypic consequences of archaic introgression and human-lineage specific biological changes.

## Materials and Methods

### Dataset

We selected 45,532 UKBB donors with available imputed genomes (Field ID 22828) and brain MRI-derived phenotypes as of October 2023 (ASEG whole brain volume, Field ID 26514, used as representative trait for filtering). Their genomes were then phased with SHAPEIT5_phase_common (*85*), on chunks of 20 cM with an overlap of 1cM and using the UKBB genotype array data as scaffolds to improve phasing. We used GLIMPSE2 (*86*, *87*) to define the sliding windows depending on the genetic map provided by SHAPEIT5. We merged the phased chunks using SHAPEIT5_ligate.

The phasing resulted in an average switch rate of 0.0036% and an average phasing quality of 98.29%, confirming accurate phasing.

Among the 45,532 UKBB samples with brain MRI-derived phenotypes described above, we selected samples identified as British with West-European ancestry (field 22006, code 1), not considered outliers for missingness and heterozygosity (field 22027), and pruned to remove kinship coefficient ≥ 0.04419 (third or lower degree relatives). This resulted in a sample size of 38,406, with 16,159,788 variants retained for having MAF in UKBB >0.001 and imputation INFO score > 0.8.

We additionally merged genomes from the 1000G dataset (generated following the pipeline in (*88*)), resulting in 13,827,267 biallelic variants. We merged the data with the three high-coverage Neanderthal genomes from Altai (*23*); http://ftp.eva.mpg.de/neandertal/altai/AltaiNeandertal/VCF/), Vindija (*24*); http://ftp.eva.mpg.de/neandertal/Vindija/VCF/Vindija33.19/) and Chagyrskaya (*25*); http://ftp.eva.mpg.de/neandertal/Chagyrskaya/VCF/) as well as the Denisova genome (*2*); http://ftp.eva.mpg.de/neandertal/Vindija/VCF/Denisova/) and the consensus human ancestral reference genome https://ftp.1000genomes.ebi.ac.uk/vol1/ftp/technical/retired_reference/ancestral_alignments/ as reconstructed by the 1000 Genomes Project consortium (*89*).

### Extraction of Neanderthal segments from the UKBB

We used SSTAR (*31*) to estimate S* score for 50-kb windows with a 10-kb step size, using the Yoruba genomes as reference and the UKBB genomes as target. S* score identifies divergent haplotypes in a target population that carry derived variants in strong LD while being absent in the reference population (*11*, *32*, *90*). We only retained segments where the observed S* score fell within the top 99^th^ percentile of a null distribution generated under a neutral model without introgression.

We generated sequences under a neutral model using SSTAR quantile and ms (*91*) following the demographic model without introgression from (*92*), with a sample size of 10,100 (--nsamp), 20,000 replicates (--nreps), diploid population size of 1,000 (--N0), target population size of 10,000 (--tgt-size), reference population size of 100 (--ref-size), sequence length of 50,000 (--seq-len), mutation rate of 1.2x10^-8^ (--mut-rate), a recombination rate of 0.7x10^-8^ per generation per base (--rec-rate) and considering 5 to 25 SNPs with a step of 5 (--snp-num-range).

We further refined the putative Neandertal segments as being delimited by Neandertal-derived variants (i.e. derived variants present in the Altai Neandertal genome and with DAF < 1% in the Yoruba genomes). We merged overlapping segments and deleted any segment with a modern human-derived variant (i.e. variants with a DAF > 1% in the Yoruba genomes and absent in the Altai Neandertal genome). We further removed segments with a match rate to Denisova 0.01 greater than that of the Altai Neandertal using SSTAR matchrate (*31*), to remove potential Denisova segments misattributed to Neandertal.

### f-statistics

We used the *f_4_-ratio* to estimate the proportion of the introgressed segments (UKBBNea) being more similar to modern humans than Neandertals. Briefly, *f_4_-ratio* represents the admixture proportion α as 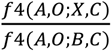, where B is the putative source of admixture, X is the putative admixed group, A is a sister group of B, C is the sister group of X and O is an outgroup to all four populations. This equation can be seen as how does the excess genetic drift between A and X (numerator) compares to the full genetic drift between A and B (denominator) (*37*). Here, we used 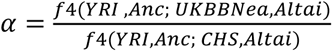, where *CHS* (represented by HG00684, a HanChinese genome) is the putative source of admixture, UKBBNea is the putative admixed population, *YRI* (represented by NA19235, a Yoruba genome) is a sister group of *CHS*, the Neanderthal *Altai* genome is the sister group of *UKBBNea*, and the ancestral genome *(Anc)* is used as the outgroup in this phylogeny. Here, we effectively compare the potential excess genetic drift between the Yoruba genome and the introgressed Neanderthal fragments (numerator) to the expected full genetic drift between the Han Chinese and the Yoruba (denominator). F-statistics were computed using POPSTATS (*93*).

### Inferring demography from the Site Frequency Spectrum using *fastsimcoal Data prepatation*

Demographic inferences were carried out using *fastsimcoal2.8* (*33, 34*) on 100 composite diploid Neanderthal introgressed genomes matching the Neanderthal introgressed haplotypes DAF spectrum, along with 100 diploid genomes from the European UKBB dataset, 87 Yoruba genomes, 100 Tuscan genomes, and the three high-coverage Neanderthal genomes. We removed positions with MAF < 0.1% in the Tuscan and Yoruba genomes to match the UKBB quality control and avoid an over-representation of rare variants which affect the SFS. We subsequently restricted the data to genomic positions that were found to be strongly affected by background selection and biased gene conversion (*35*, *94*). Briefly, we kept only positions for which the ancestral state is known, positions with a recombination rate >1 cM/Mb using the HapMap Phase II genetic map on b37 (https://github.com/odelaneau/shapeit4/tree/master/maps/) (*95*), outside of CpG islands using the UCSC genome annotation on GRCh37 (https://hgdownload.soe.ucsc.edu/goldenPath/hg19/database/cpgIslandExtUnmasked.txt.gz) (32), not a CpG site, and with A↔T and G↔C mutations which are not affected by biased gene conversion. We retained 283,506 polymorphic positions. We estimated a new average recombination rate on the neutral set of positions to 5.88x10^-8^ recombination.bp^-1^.generation^-1^. The VCF was converted into a joined unfolded SFS using easySFS (*96*).

### Fastsimcoal simulations

Because introgressed Neanderthal haplotypes in European UKBB individuals are shaped by post-introgression European demography, we first estimated demographic parameters for the three high-coverage Neanderthal genomes, European individuals from UKBB, and Yoruba, without including the introgressed Neanderthal. We specified the inbreeding coefficient (FIS) to be 0.162 for Altai, 0.062 for Vindija and 0.193 for Chagyrskaya, estimated based on the proportion of homozygous-by-descent track >2.5 cM using a constant recombination rate inferred in (*25*). We also replicated the demography simulations using the TSI genomes from the 1000 genome project instead of the European UKBB to test for the accuracy of the estimates and the robustness towards data generation differences.

We next incorporated an explicit introgressed Neanderthal lineage into the *fastsimcoal* demographic model to infer its divergence time (T_UKBB-Vindija_divergence_), effective population size (N_UKBBNea_pre-intro_), and bottleneck associated with introgression (N_UKBBNea_intro_, _TUKBBNea_end_intro_, T_UKBBNea_end_intro_). We used the growth rate *G_UKBB_* and the onset time of growth *T_UKBB_growth_* (Table S1)ーto model the post-admixture evolution of the introgressed Neanderthal segments. However, we allowed the exact effective population sizes pre- and post-growth (N_UKBBNea_postintro_ and N_UKBBNea_, Table S5) to be freely inferred in *fastsimcoal*. Potential biases in our introgression detection methodology—including over-assignment of fragments matching Altai-specific variants and a low rate of false-positive introgressed segments that are actually modern humans—can bias our inference on the UKBB Neanderthal demography. We circumvent this issue by allowing migrations from both Altai and UKBB into the introgressed lineage to account for the potential excess affinity towards these two groups.

To validate the relationships among the three high-coverage Neanderthal genomes and the introgressed lineage, and to inform model specification, we used allele frequency–based approaches *TREEMIX* (*36*) and *qpGraph* (*37*). We generated trees with zero to five migration edge(s) with *TREEMIX*, on the same SNPs as used in fastsimcoal simulations (n=283,506 SNPs), setting YRI as the root and using 20 SNPs per block for estimation of the covariance matrix. We generated fully automated qpGraphs using the find_graph() function of ADMIXTOOLS 2.0.4 R package (*97*), inferring zero to three admixture events on the composite UKBB Neanderthal used in *fastsimcoal* simulations, the high-coverage Neanderthals, YRI and the ancestral genome used as root. This analysis was computed on the intersection between the 283,506 SNPs used in the *fastsimcoal* simulations and the ancestral genome, resulting in 202,574 SNPs. For each number of admixture events, the best *qpGraph* was selected over 20 simulations.

*fastsimcoal2.8* was performed on 200,000 coalescent simulations (-n 200000) to estimate the expected derived SFS (-d) using 50 optimisation (ECM) cycles (-L 50) and with parameter estimation (-M). The best parameter estimates were selected based on the maximum likelihood over 100 independent runs. Because an accurate mutation rate for the neutral SNP set was unavailable, we fixed the effective population size of the Yoruba at t_0_=28,500, based on previous estimates obtained using SFS methods (*98*).

Confidence intervals were estimated using a parametric block-bootstrapping approach, in which we first generated 20 SFS by randomly sampling 100 blocks of 2,835 SNPs. For each of the 20 bootstrapped SFS, we re-estimated the model parameters using 20 independent runs on the aforementioned coalescence simulations. The 95% confidence intervals were estimated on the distribution of the 20 newly estimated ML parameter values.

Template and estimation files specifying the demographic models, and the parameter estimation of the maximum likelihood simulations are provided in Tables S1-S5.

### Genome-wide detection of introgressed Neanderthal variants deviating from Hardy-Weinberg equilibrium

#### Hardy-Weinberg equilibrium exact-test on introgressed haplotypes

The Hardy-Weinberg equilibrium exact test was applied to a modified VCF in which introgressed haplotypes were encoded as the ‘1’ allele and non-introgressed haplotypes as the ‘0’ allele. This recording allowed us to identify genomic positions exhibiting an excess of introgressed haplotype relative to expectation under the Hardy-Weinberg equilibrium, given the overall introgression frequency. Hardy-Weinberg statistics were computed using plink2 --hardy (*99*, *100*).

We defined candidate loci as sites showing a significant excess of Neanderthal-introgressed homozygote, with an HWE exact-test p-value < 10^-8^, and a count of observed introgressed homozygotes greater than the expected count under equilibrium. To ensure that an excess of introgressed homozygotes specifically drove the deviation from HWE, we further required that (i) the deviation of the introgressed homozygotes from their expected number exceed that of non-introgressed homozygotes, and (ii) the number of observed heterozygotes was lower than expected. We additionally restricted detection to variants with an introgression rate of 0.5% or less to control for false positives in the introgression detection. We focused on alleles of Neanderthal origin, and removed alleles that likely arose in the common ancestor of both modern humans and Neanderthals, or that arose after introgression specifically in the European lineage. To do so, we first restricted our analysis to variants matching the Altai, Vindija, or Chagyrskaya Neanderthal genomes and absent from the Yoruba population (346,401 variants). Then, we extracted variants that are likely of Neanderthal origin but not observed in any of the sequenced Neanderthals. Because Neanderthal-derived alleles are expected to be tightly linked to introgressed haplotypes, their DAF on these haplotypes should be substantially higher than that of variants arising post-introgression. We therefore estimated the DAF distribution of sequenced Neanderthal-specific alleles on introgressed haplotypes and used this distribution to infer additional candidate Neanderthal-private variants that were not observed in the three sequenced Neanderthal genomes. Specifically, we retained variants whose DAF on the introgressed haplotype fell within the upper 95% of the Neanderthal-specific distribution (DAF > 0.69, Fig. 3a). Using this strategy, we identified 50,680 derived variants absent from the Yoruba population and from all sequenced Neanderthals, yet strongly associated with introgressed segments and exhibiting high frequency on Neanderthal haplotypes, consistent with a Neanderthal origin. We refer to this new set of 397,081 variants as Neanderthal-specific variants.

#### Functional annotation of excess Neanderthal-introgressed homozygotes

We clumped candidate sites showing an excess of Neanderthal-introgressed homozygotes with PLINK 1.9 (--clump --clump-r2 0.2 --clump-kb 100) (*99*, *101*). Within each linkage disequilibrium (LD) block (r2 > 0.2 within 100-kb), we retained the variant with the lowest HWE exact-test p-value. To exclude sites in strong HWE deviation caused by the hitchhiking effect of a nearby non-introgressed site, we excluded sites for which a non-introgressed site in LD (r^2^>0.2 within 100 kb) displayed a more significant HWE deviation. Sites in LD were recovered using PLINK v2.0 (--r2-phased --ld-window-r2 0.2 --ld-window-kb 100) (*99*, *100*).

We tested for Gene Ontology (GO) enrichment using GREAT v4.0.4 (*102*). Each clumped site was associated with nearby genes using the default “basal plus extension” model, in which each gene is assigned a basal regulatory domain spanning 5 kb upstream and 1 kb downstream of the transcription start site, irrespective of neighbouring genes. These basal domains are then extended bidirectionally to the nearest gene’s basal domain, up to a maximum of 1,000 kb. We tested enrichment for biological terms relative to a genomic background matched for DAF and LD score to the clumped Neanderthal introgressed, to avoid bias arising from the non-random DAF and LD-score distribution of these specific Neanderthal variants (Wilcoxon test, p < 10^-15^ for both, Fig. S8). This correction is motivated by the observation that both MAF and LD-score correlate with distance to genes (Spearman’s correlation test, r = 0.02, p < 10^-15^ and r = 0.04, p < 10^-15^, respectively, Figs. S9 and S10), and therefore with the power to detect GO enrichments. LD-score was computed as the sum of r^2^ values across 200-kb windows genome-wide using PLINK v1.9 (--r2 --ld-window-kb 200 --ld-window-r2 0) (*99*, *101*). Background sets were matched to the target variants using 5%-wide DAF bins and LD-score bins of width 2. We generated 1,000 matched background replicates for enrichment testing.

We further annotated the clumped sites with CADD v1.17 (*103*) (https://kircherlab.bihealth.org/download/CADD/v1.7/GRCh37/), the NHGRI-EBI Catalogue of human genome-wide association studies v1.0.2 (downloaded on 03/12/2025) (*56*) and expression Quantitative Trait Loci (eQTL) from GTEx v8 (*57*) (https://ftp.ebi.ac.uk/pub/databases/spot/eQTL/imported/GTEx_V8/ge/, downloaded on 26/01/2026). Genomic coordinates from the GWAS catalogue and GTEx dataset were converted from hg38 to hg19 using liftOver (*104*). We tested whether sites showing an excess of Neanderthal-introgressed homozygotes were enriched for suggestive GWAS associations (p < 1×10⁻^5^), keeping only the strongest hit for each position. We likewise tested whether these sites were enriched for suggestive eQTLs (p < 1×10⁻^5^) in each GTEx tissue, using a Bonferroni-corrected significance threshold of 0.05/49. For both analyses, we used the same DAF- and LD-score–matched genomic background as in the GREAT enrichment, because minor allele frequency and LD structure are known to influence the power to detect significant associations (*53–55*). The clumped set were additionally annotated with the regions identified ad adaptive introgression in (*49*) and with F_ST_ between the GBR and YRI in the 1000G dataset computed with PLINK v 2.0 (*99*, *100*) using the command --fst and using as filter --geno 0.01, as well as their genome-wide percentile category estimated with the percent_rank() function of the dplyr R package (*105*).

### Detecting Neanderthal introgression depletion

To define Neanderthal gaps, we first selected 10-kb windows in which the mean inferred Neanderthal introgression rate was < 10⁻³ and in which no site had an introgression probability ≥ 10⁻². A window size of 10 kb was chosen because it is smaller than the estimated average length of introgressed Neanderthal segments, ensuring that introgressed tracts are not systematically diluted by windowing. Among windows with low inferred introgression, we further distinguished two categories:

i. windows containing at least one Neanderthal-specific SNP, defined as a site carrying the Altai Neanderthal allele and with frequency <1% in Yoruba individuals, referred to as ‘strict gaps’; and
ii. windows containing no Neanderthal-specific SNPs, which likely reflect ILS. This distinction reflects the fact that windows lacking Neanderthal-specific variants are undifferentiated between Yoruba and Neanderthals and therefore have limited power to detect introgression. Such windows cannot be confidently interpreted as being depleted of Neanderthal ancestry. To quantify the expected frequency of undifferentiated windows, we estimated the number of mutations private to the Altai lineage relative to Yoruba as μ × T × L = 1.2 × 10⁻⁸ × 20,200 × 10,000 = 2.4, where μ is the mutation rate per base per generation, *T* is the divergence time in generations, and *L* is the window length. Thus, even in a 10-kb window, a substantial fraction of regions is expected to lack informative Neanderthal-specific polymorphisms by chance alone. All genomes were masked using the pilot accessibility mask (ftp://ftp.1000genomes.ebi.ac.uk/vol1/ftp/release/20130502/supporting/accessible_genome_masks/20141020.pilot_mask.whole_genome.bed, (*89*)) to remove regions of low mappability, which are prone to genotyping error, particularly in the Altai Neanderthal genome.

Finally, continuous ILS windows were assigned to gaps if they were flanked by strict gaps on both sides. We refer to this set of strict gaps and newly assigned gaps as expanded gaps. Likewise, we assigned continuous ILS windows to introgressed windows if they were flanked by introgressed windows on both sides, and refer to them as expanded introgressed windows.

All 10-kb windows were annotated with several measures of constraints and protein interaction, corrected for the proportion of overlap between the 10-kb window and the gene. We compared the mean of Sum Singleton Cohort (SSC) score (*106*), number of protein-protein interactions (PPI) (*107*), intolerance to variation (RVIS) (*108*), probability of loss of function (pLI), sensitivity to missense Z-score (mis_Z), and sensitivity to loss-of-function Z-score (Lof_Z) (*109*) between the three different categories using pairwise Wilcoxon tests implemented in R v.4.4.3 (*110*). The windows were also annotated with their average recombination rate using the HapMap Phase II genetic map on b37 (https://github.com/odelaneau/shapeit4/tree/master/maps/) (*95*).

We tested which 10-Mb regions are most depleted in Neanderthal introgression using our expanded gap set and compare them to the four previously identified long introgression deserts (chr1:105,000,000-121,000,000; chr3:74,100,000-89,300,000; chr7:106,000,000- 123,000,000; chr8:49,400,000-66,500,000, (*10*)). We scanned the genome in 10-Mb windows with a step size of 1,000,000 bp and calculated the proportion of expanded gaps in each windows. Windows were then classified by percentile, and the 1 and 5% most depleted were extracted. Adjacent depleted windows were merged using bedtoolsr R package (*111*, *112*) computing the average desertness of the top windows. We assessed the overlap with the previously identified 10-Mb long desert.

We additionally extracted the 0.1% longest continuous stretch of expanded gaps, detect the overlapping genes using biomaRt R package (*113*) and Ensembl 115 (*114*). We run a network analysis using the online software STRING (https://string-db.org/v12.0) looking for the extent of protein-protein interactions formed by this gene list. We also performed a functional enrichment analysis using the same software and highlighted the genes leading the reported enrichment.

The expanded Neanderthal gaps were then annotated with HARs (*68*), 50-kb region TMRCA within the Yoruba, between modern human and Altai Neanderthal, and between modern human and Denisova using the method described in (*67*), the NHGRI-EBI Catalogue of human genome-wide association studies v1.0.2 (downloaded on 03/12/2025) (*56*) and expression eQTLs from GTEx v8 (*57*) (https://ftp.ebi.ac.uk/pub/databases/spot/eQTL/imported/GTEx_V8/ge/, downloaded on 26/01/2026). We tested whether genome categories (gap, introgressed ILS) were enriched in HARs, comparing the point estimate to random genomic segments similar in size and length to the HARs set, using 1,000 permutations. The same approach was done within each of the four TMRCA categories. Enrichments of NHGRI-EBI GWAS catalog and GTEx hits on the expanded deserts was tested against a DAF and LD-score matching genomic background of 1000 replicates, as described in the above section, after LD-pruning of the expanded gaps using PLINK v1.9 (*99*, *101*) --indep-pairwise 100 10 0.2. The p-values were corrected using the Benjamini-Yekutieli method from the R package sgof (*115*).

## Acknowledgements

The analyses in this paper were performed using the resources and support provided by University of Padova Strategic Research Infrastructure Grant 2017: “CAPRI: Calcolo ad Alte Prestazioni per la Ricerca e l’Innovazione” and by the BioData Hub at the Department of Biology, University of Padova. The BioData Hub was established with funding from the Italian Ministry of University and Research (MUR) ‘Departments of Excellence’ program 2023-2027, as part of the project “Biological networks: from molecules to ecosystems.” This research was funded by the European Union - Next Generation EU, Mission 4 Component 1, CUP D53D23016690001. This research has been conducted using the UK Biobank Resource under application number 86275. We thank Jens Blöcher (Johannes Gutenberg-Universität Mainz) for helpful suggestions on fastsimcoal simulations.

