## Supplementary Figures for "The Neanderthal population history and the introgression landscape inferred from the UK Biobank"

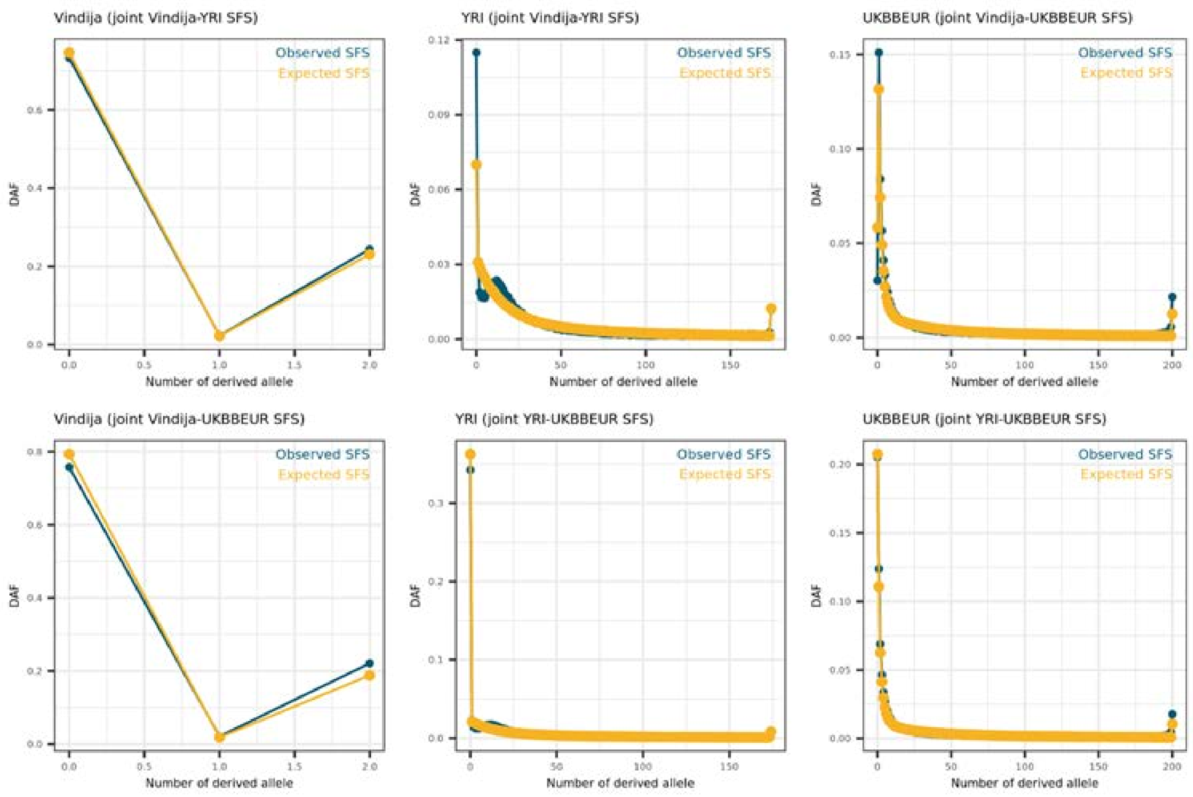


**Fig S1. SFS fit of the modern genomes Maximum Likelihood (ML) simulation with post-Out of Africa growth.** Observed (blue) and expected (yellow) derived allele distribution of the modern genomes demography Maximum Likelihood simulation (European UKBB and Yoruba), considering an eventual population growth after the Out of Africa. The maximum observed likelihood is -1,636,753.462. This model resulted in the smallest difference between the estimated maximum likelihood and observed maximum likelihood (Δ = 44,280.233), compared with the model where population size remained constant (Δ = 53,050.317; Fig S2). The specific modelling specification (template and estimate files) and the ML estimated parameters are described in Table S1.

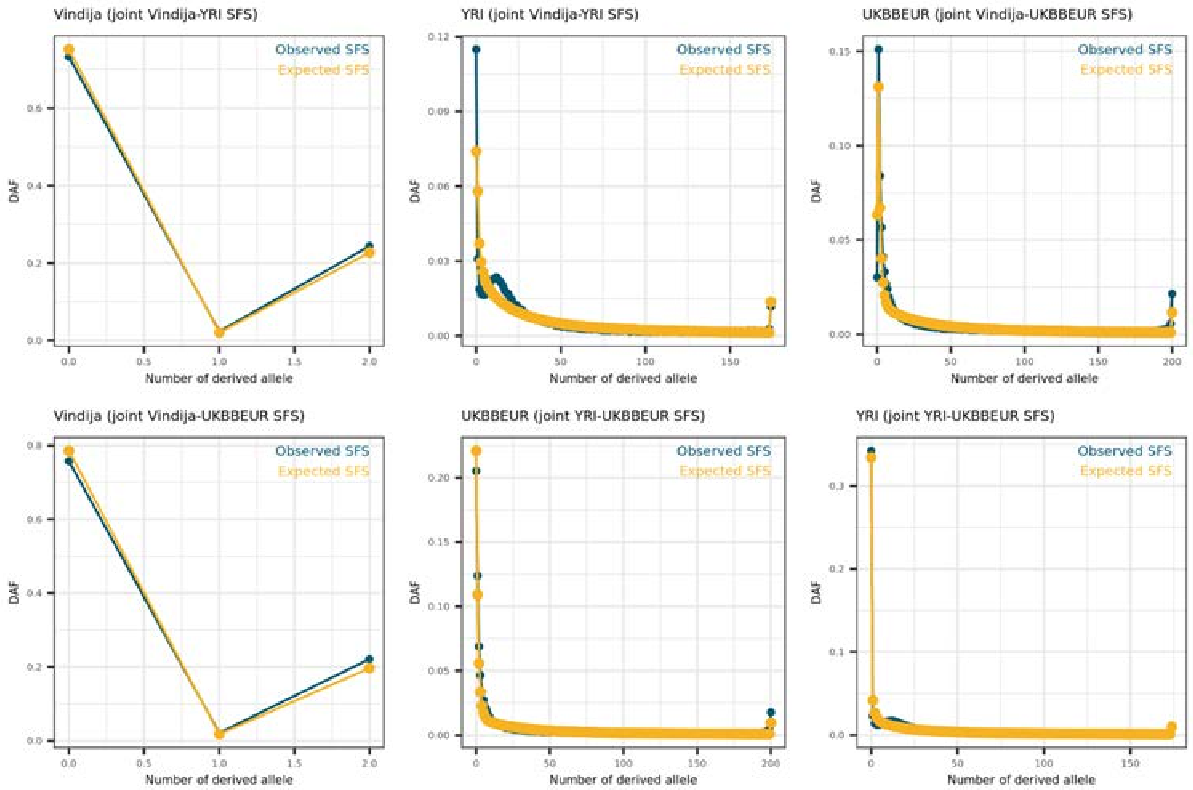


**Fig S2. SFS fit of the modern genomes Maximum Likelihood simulation without post-Out of Africa growth.** Observed (blue) and expected (yellow) derived allele distribution of the modern genomes demography Maximum Likelihood simulation (European UKBB and Yoruba), without any population growth after the Out of Africa. The maximum observed likelihood is -1,636,753.462, and the difference between observed expected and observed likelihood Δ = 53,050.317. The specific modelling specification (template and estimate files) and the ML estimated parameters are described in Table S2.

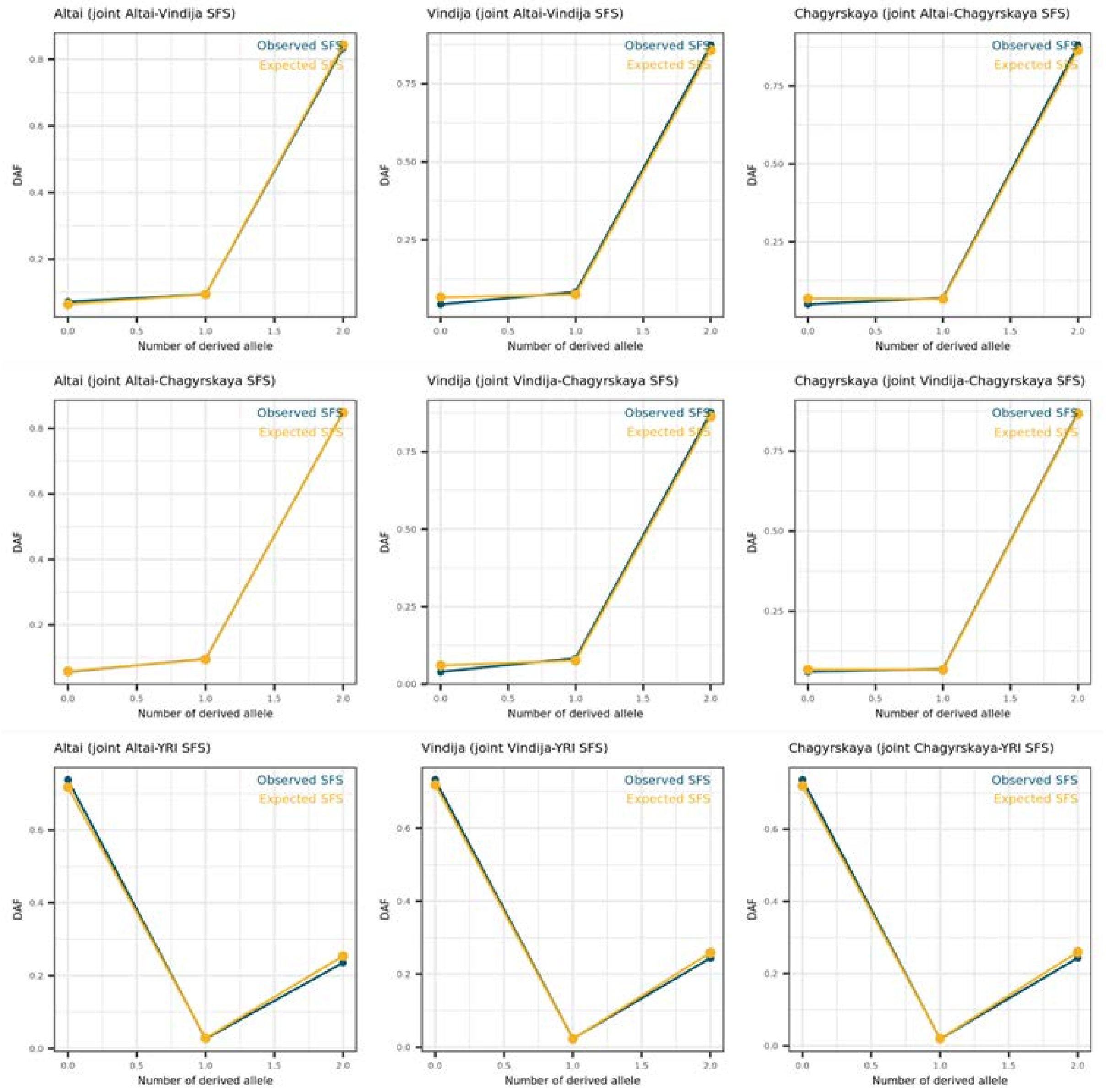


**Fig S3. SFS fit of the archaic genome Maximum Likelihood simulation.** Observed (blue) and expected (yellow) derived allele distribution of the archaic genomes demography Maximum Likelihood simulation (high-coverage Neanderthals). The maximum observed likelihood is -1,293,156.226 and the difference between expected and observed likelihood Δ = 36,748.141. The specific modelling specification (template and estimate files) and the ML estimated parameters are described in Table S3.

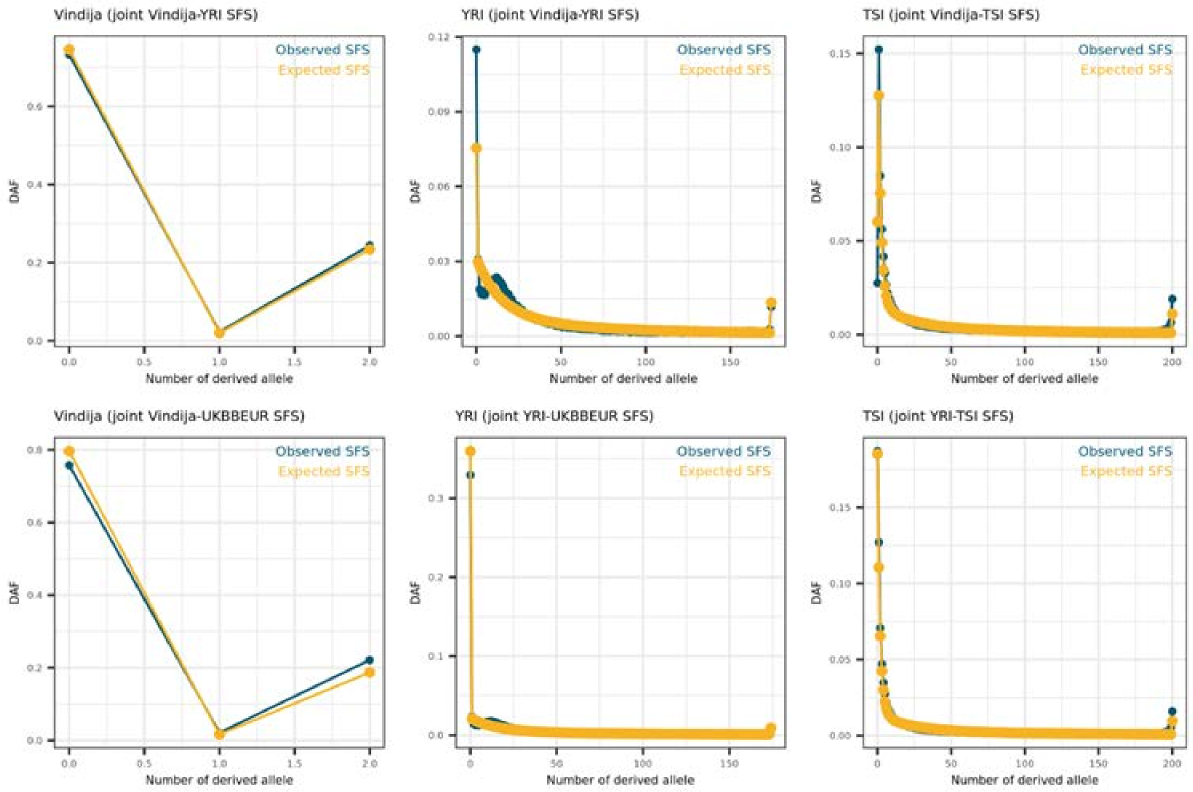


**Fig S4. SFS fit of the modern genomes Maximum Likelihood simulation with post-Out of Africa growth, using TSI.** Observed (blue) and expected (yellow) derived allele distribution of the modern genomes demography Maximum Likelihood simulation (TSI and Yoruba), considering an eventual population growth after the Out of Africa. The maximum observed likelihood is – 1,619,295.843 and the difference between expected and observed likelihood Δ = 41,020.705. The specific modelling specification (template and estimate files) and the ML estimated parameters are described in Table S4.

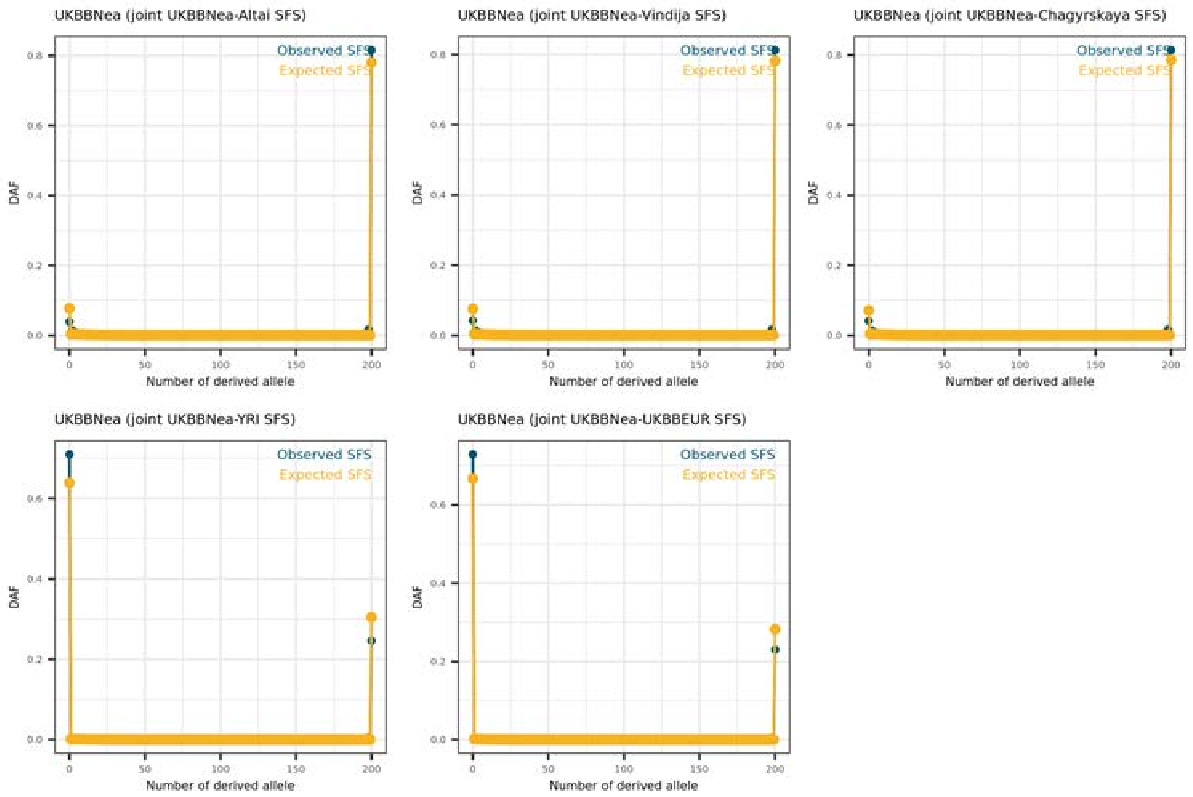


**Fig S5. SFS fit of the introgressed Neanderthal Maximum Likelihood simulation.** Observed (blue) and expected (yellow) derived allele distribution of the modern genomes demography Maximum Likelihood simulation (TSI and Yoruba), considering an eventual population growth after the Out of Africa. The maximum observed likelihood is –4,437,442.327 and the difference between expected and observed likelihood Δ = 165,887.713. The specific modelling specification (template and estimate files) and the ML estimated parameters are described in Table S5.

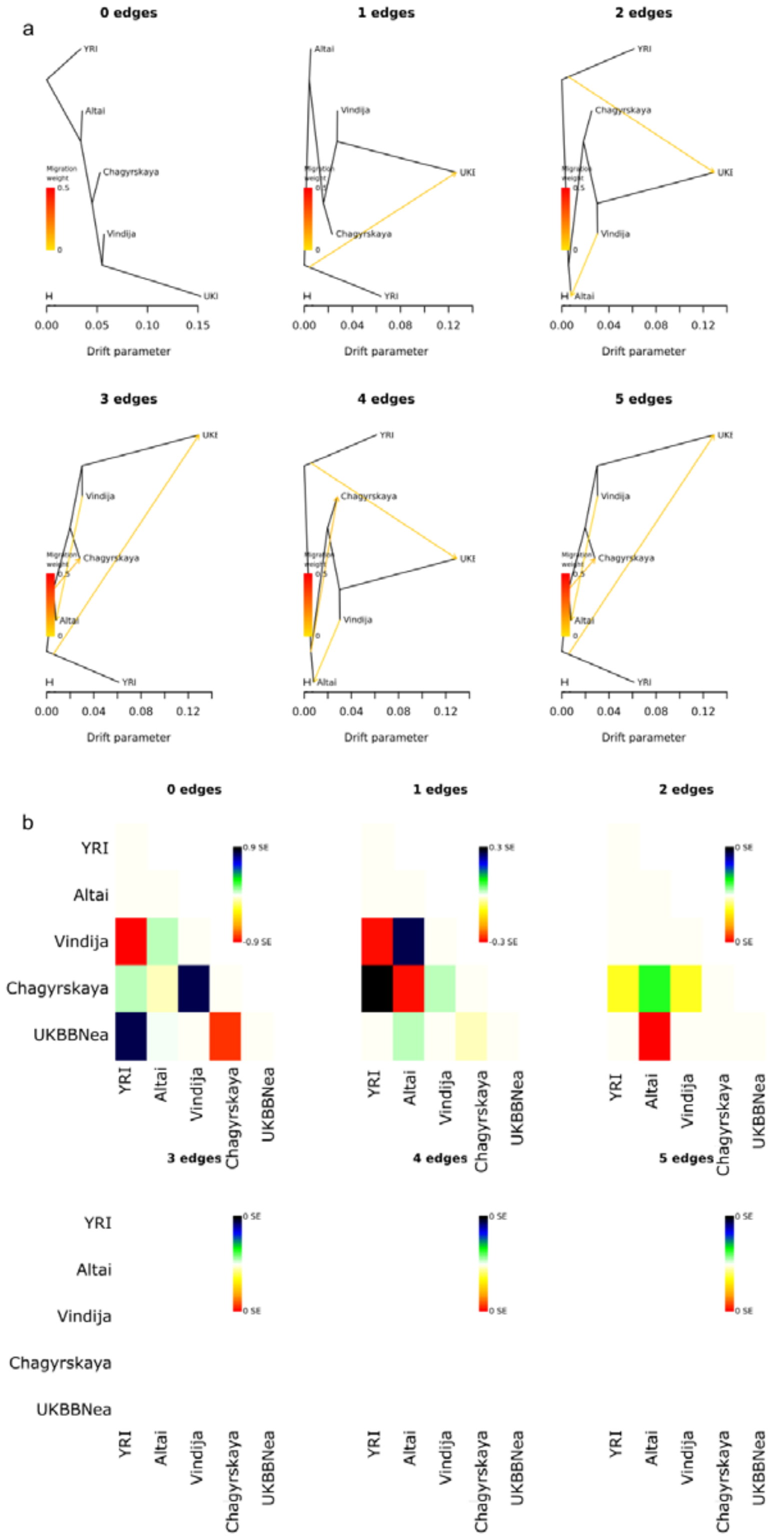


**Fig S6. Neanderthals' population split and mixture using *TREEMIX*.** Trees **(a)** and residuals **(b)** of *TREEMIX* inferences using 1 to 5 edges on the 283,506 neutral SNPs covered in all high-coverage Neanderthals and introgressed in the European UKBB, used to infer *fastsimcoal* simulations.

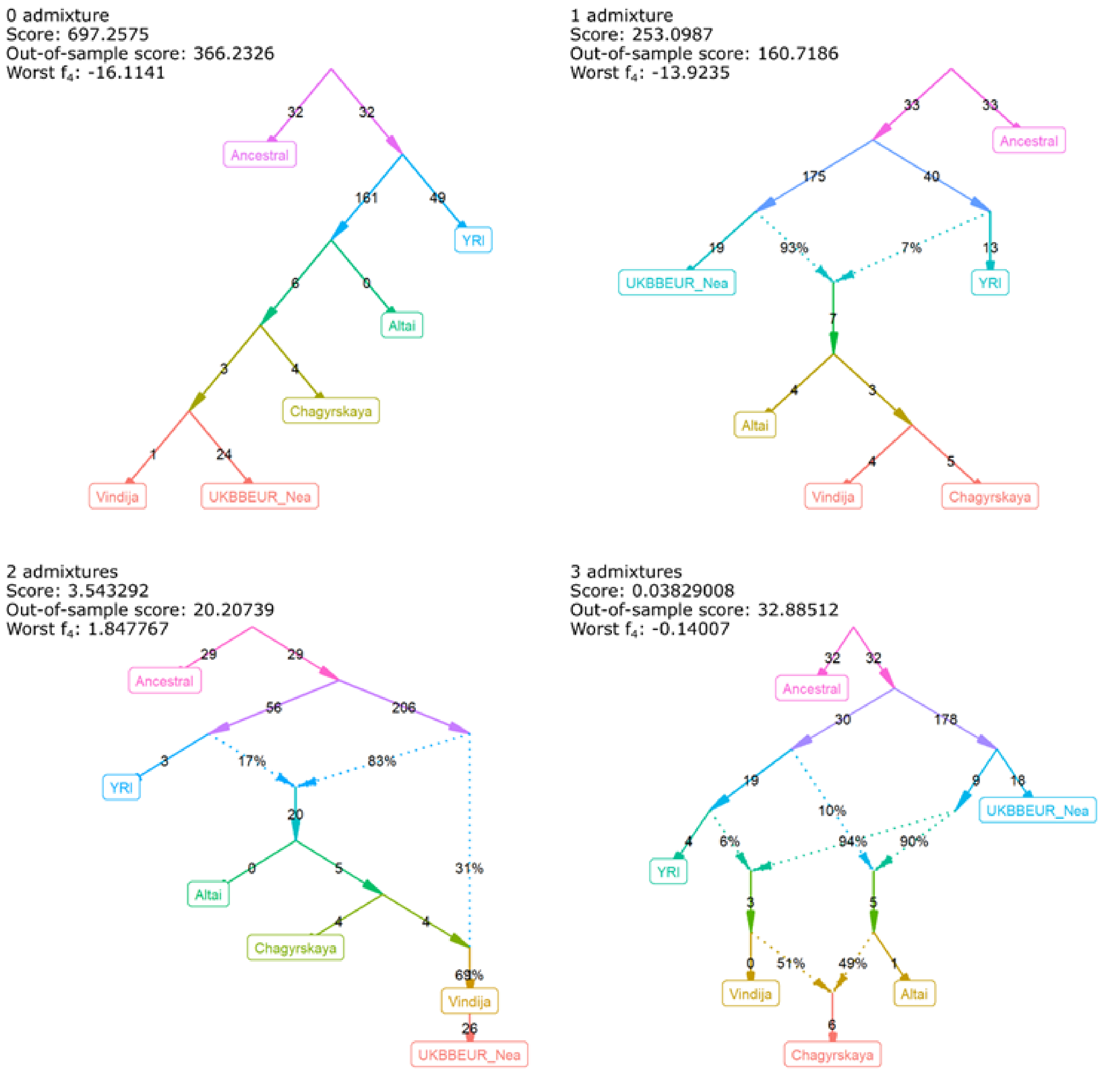


**Fig S7. Neanderthals' population split and mixture using *qpGraph*.** Fully automated graphs were used to generate trees with zero to three admixture events over 20 iterations each. We consider the two-admixture event graph as best-fitting the data based on the lowest out-of-sample score estimates. The graphs were generated on the intersection of the 283,506 neutral SNPs covered in the composite introgressed Neanderthal in the European UKBB, the high-coverage Neanderthal and the YRI genomes, used to infer *fastsimcoal* simulations, and the ancestral genome, resulting in 202,574 SNPs.

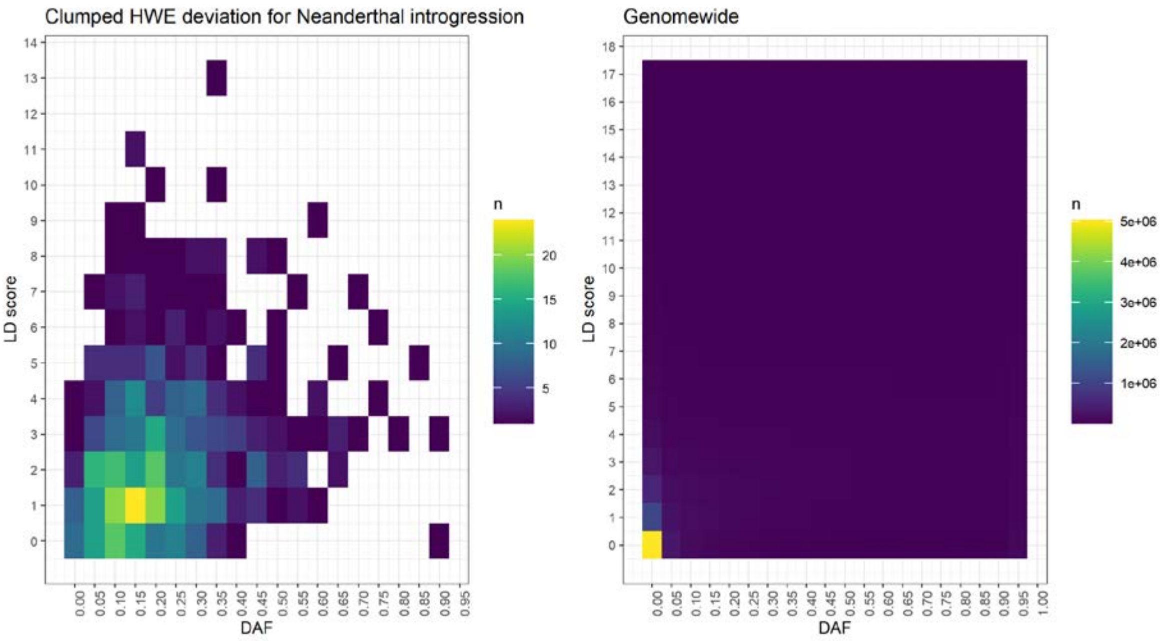


**Fig S8. DAF and LD-score distribution between the 545 clumped Neanderthal-specific variants in Hardy-Weinberg Equilibrium (HWE) deviation (a) compared to the genome average (b).**

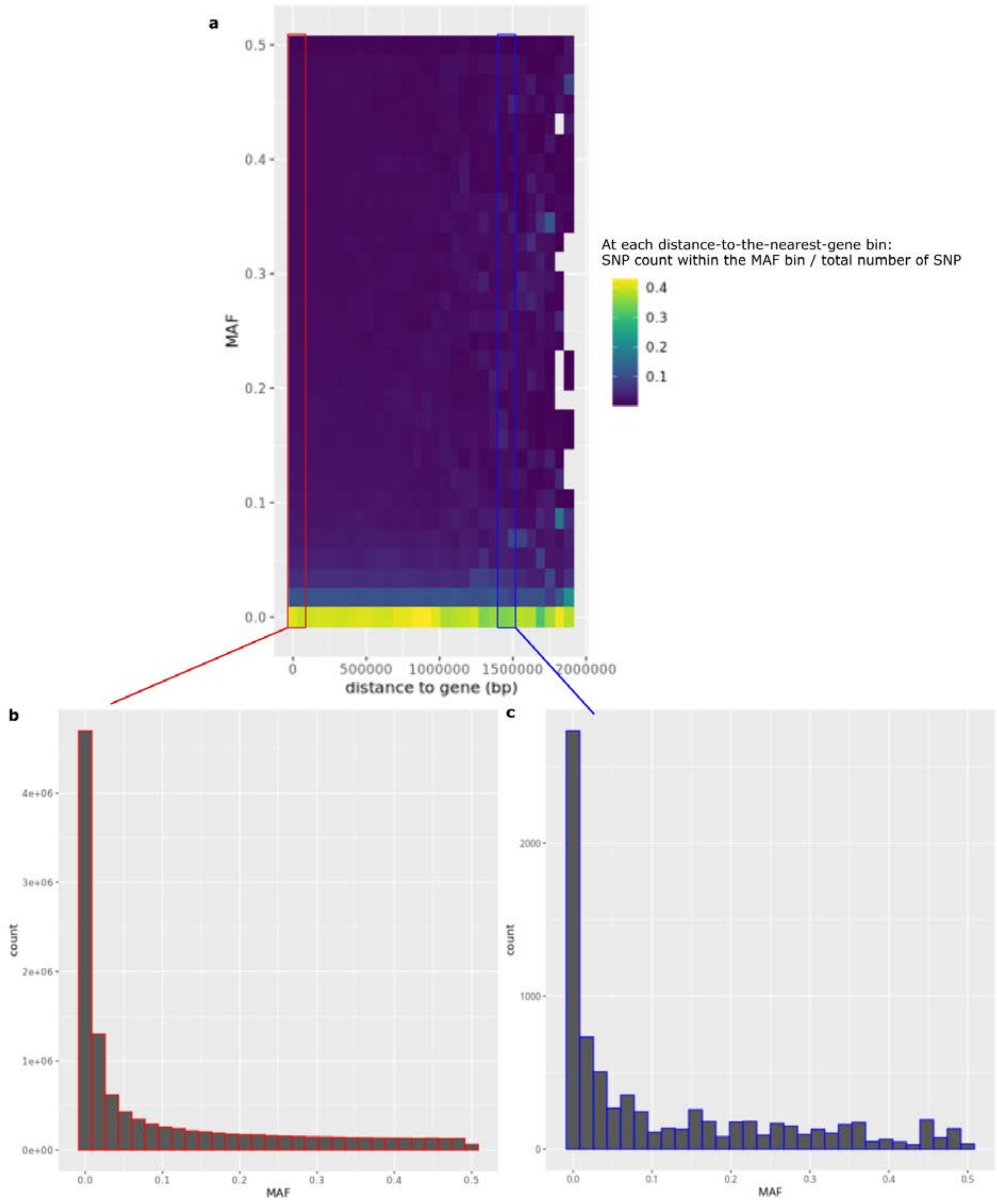


**Fig S9. Relationship between MAF and distance to the nearest gene. (a)** Distribution of the fraction of SNPs within each MAF bin (y-axis) across distance-to-the-nearest-gene bins (x-axis). As the distance from genes increases, the relative fraction of SNPs in higher MAF bins increases. **(b)** MAF distribution of SNPs located 0-100 kb from a gene, within the red box in (a). **(c)** MAF distribution of SNPs located 1,400-1,500 kb from a gene, within the blue box in (a). Compared with SNPs near genes (0–100 kb), SNPs located further away (1,400–1,500 kb) show a relative enrichment of variants in higher MAF bins.

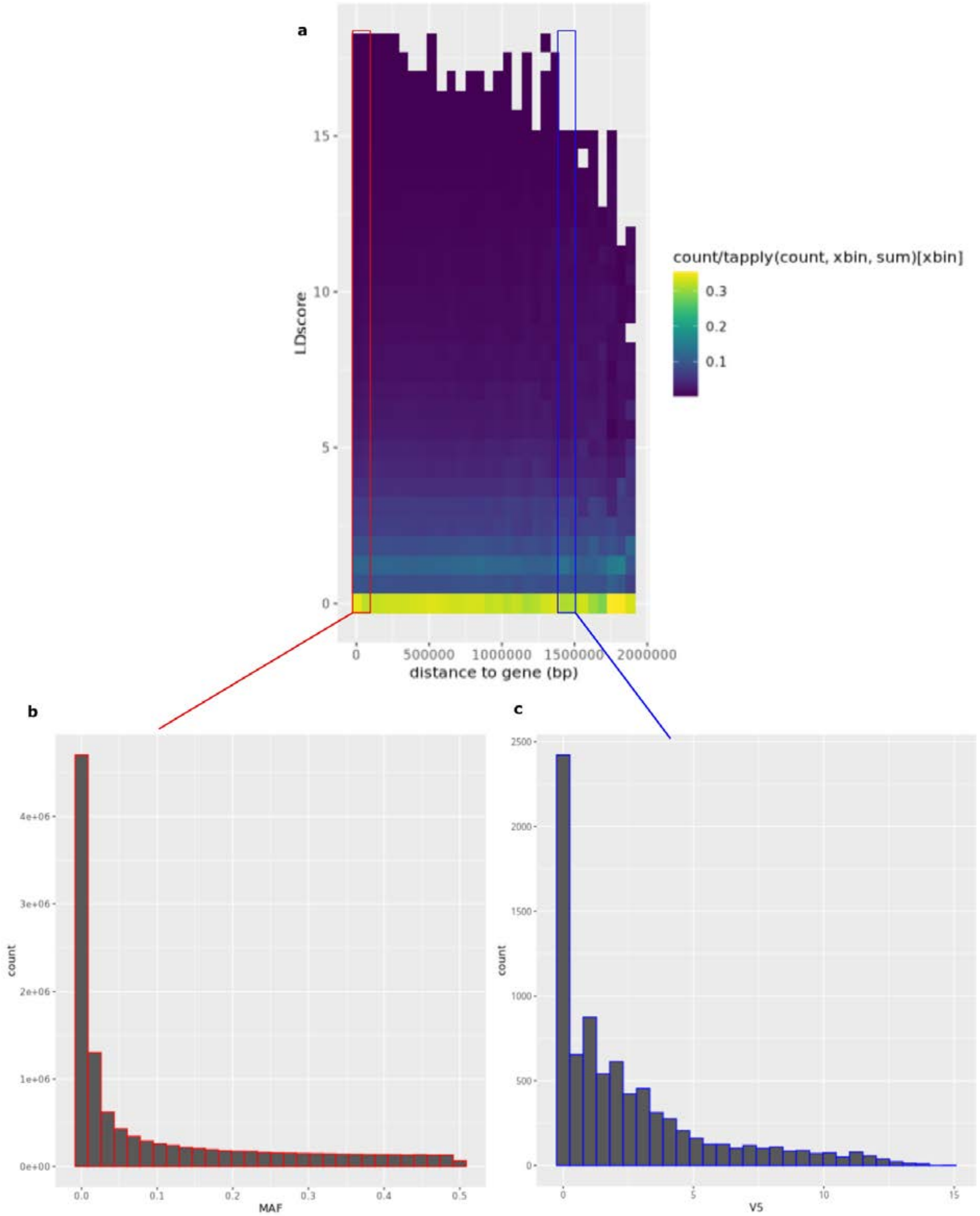


**Fig S10. Relationship between LD-score and distance to genes. (a)** Distribution of the fraction of SNPs within each LD-score bin (y-axis) across distance-to-a-gene bins (x-axis). As the distance from genes increases, the relative fraction of SNPs in higher LD-score bins increases. **(b)** LD-score distribution of SNPs located 0-100 kb from a gene. **(c)** LD-score distribution of SNPs located 1,400-1,500 kb from a gene. Compared with SNPs near genes (0–100 kb), SNPs located further away (1,400–1,500 kb) show a relative enrichment of variants in higher LD-score bins.

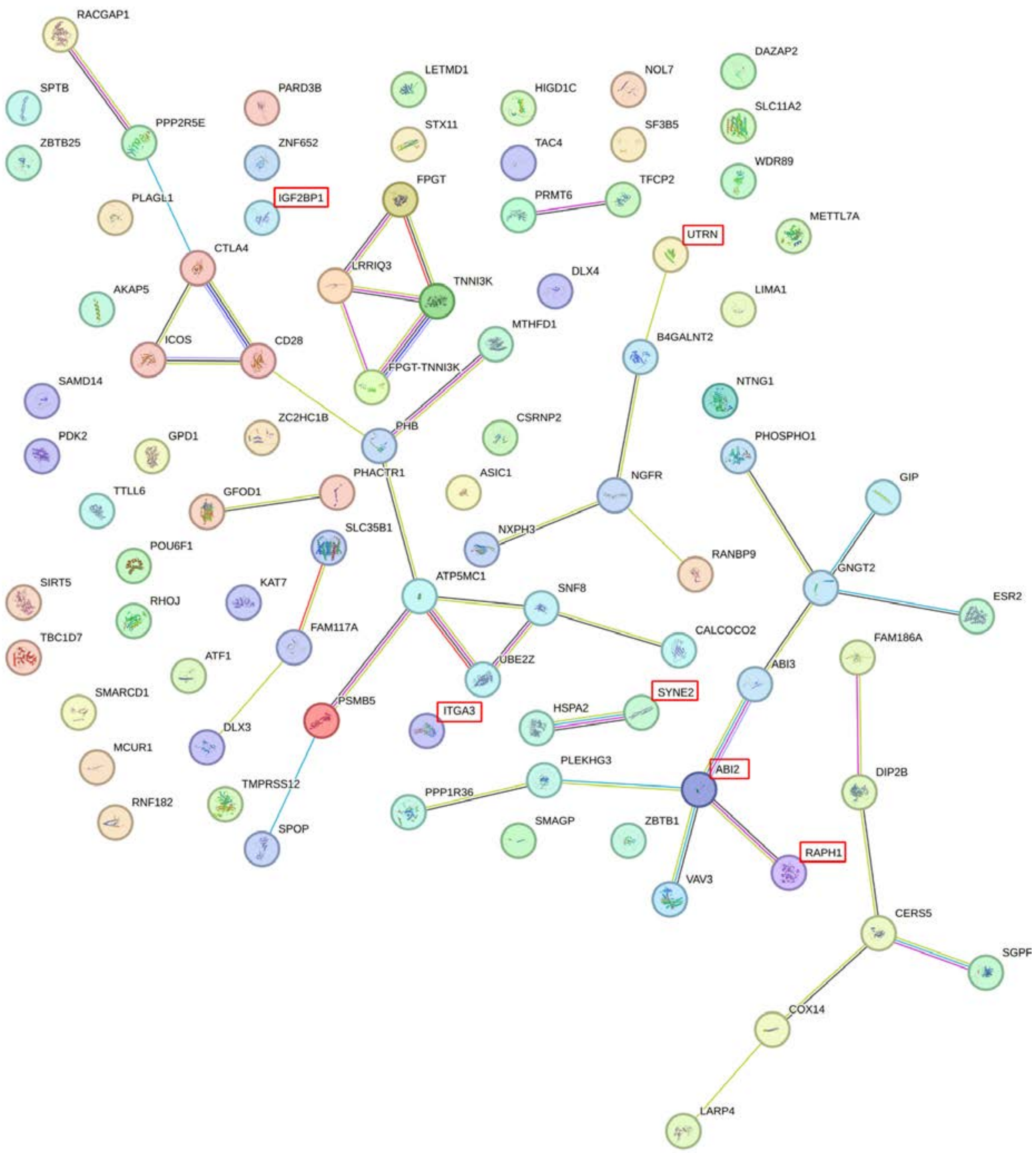


**Fig S11. Protein-Protein Interaction Network Analysis of the 98 genes in the top 0.1% longest expanded gaps.** The six genes involved in the significant filopodium-associated GO term (GO:0030175, FDR-adjusted p = 0.0215) are highlighted in the red squares. The graph was obtained using STRING v12.0 online at https://string-db.org/

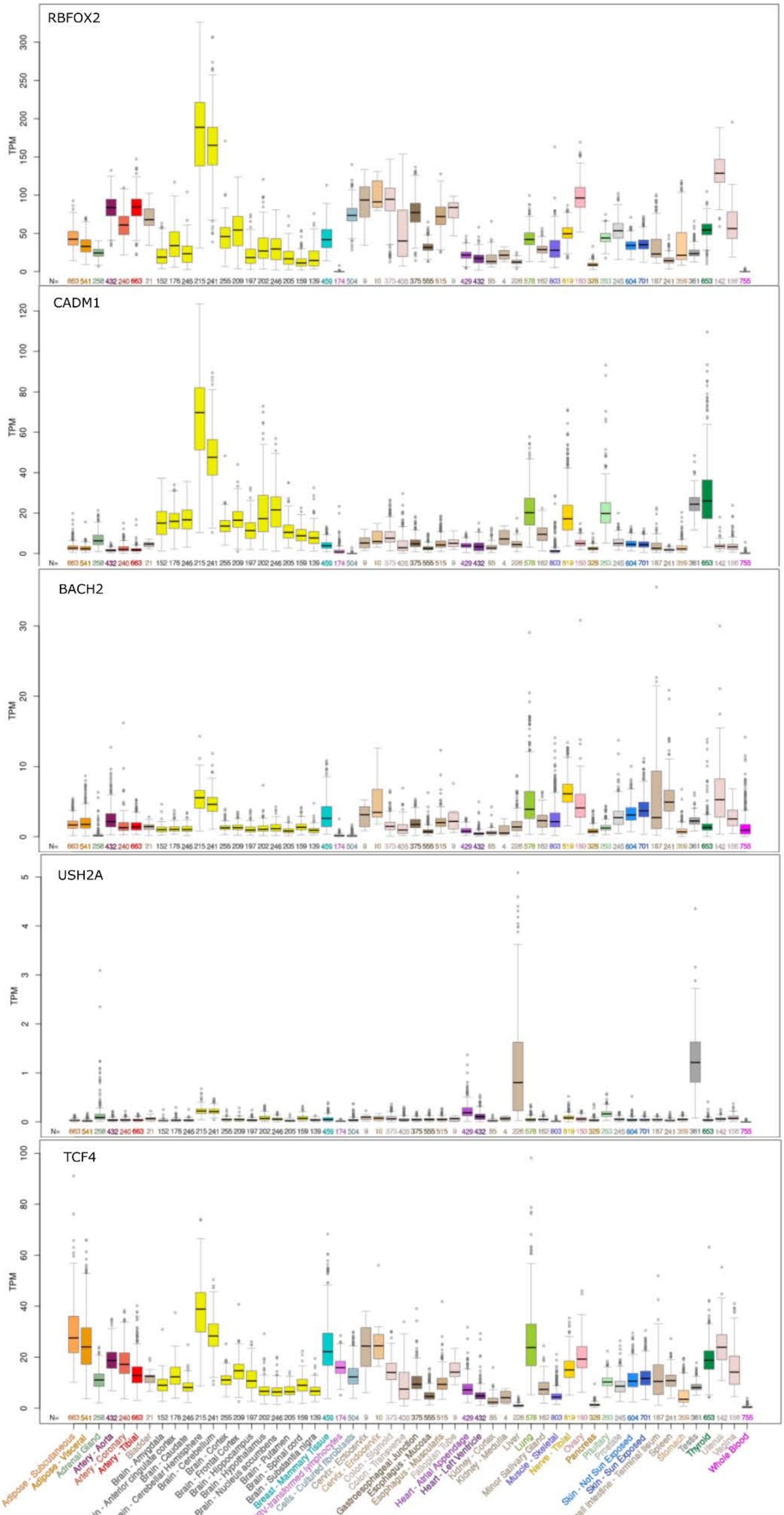
RBFOX2
CADM1
BACH2
USH2A
TCF4
TPM
N=
Adipose - Subcutaneous
Adipose - Visceral
Adrenal Gland
Artery - Aorta
Artery - Coronary
Artery - Tibial
Bladder
Brain - Amygdala
Brain - Anterior cingulate cortex
Brain - Caudate
Brain - Cerebellar Hemisphere
Brain - Cerebellum
Brain - Cortex
Brain - Frontal Cortex
Brain - Hippocampus
Brain - Hypothalamus
Brain - Nucleus accumbens
Brain - Putamen
Brain - Spinal cord
Brain - Substantia nigra
Breast - Mammary Tissue
Cells - EBV-transformed lymphocytes
Cells - Cultured fibroblasts
Cervix - Ectocervix
Cervix - Endocervix
Colon - Sigmoid
Colon - Transverse
Esophagus - Gastroesophageal Junction
Esophagus - Mucosa
Esophagus - Muscularis
Fallopian Tube
Heart - Atrial Appendage
Heart - Left Ventricle
Kidney - Cortex
Kidney - Medulla
Liver
Lung
Minor Salivary Gland
Muscle - Skeletal
Nerve - Tibial
Ovary
Pancreas
Pituitary
Prostate
Skin - Not Sun Exposed
Skin - Sun Exposed
Small Intestine - Terminal Ileum
Spleen
Stomach
Testis
Thyroid
Uterus
Vagina
Whole Blood

**Fig S12. UCSC GTEx expression profile for the six genes overlapping HARs and extended gaps with a recent modern human TMRCA (< 650 kya), and a deep modern human-Neanderthal TMRCA (≥ 650 kya).**

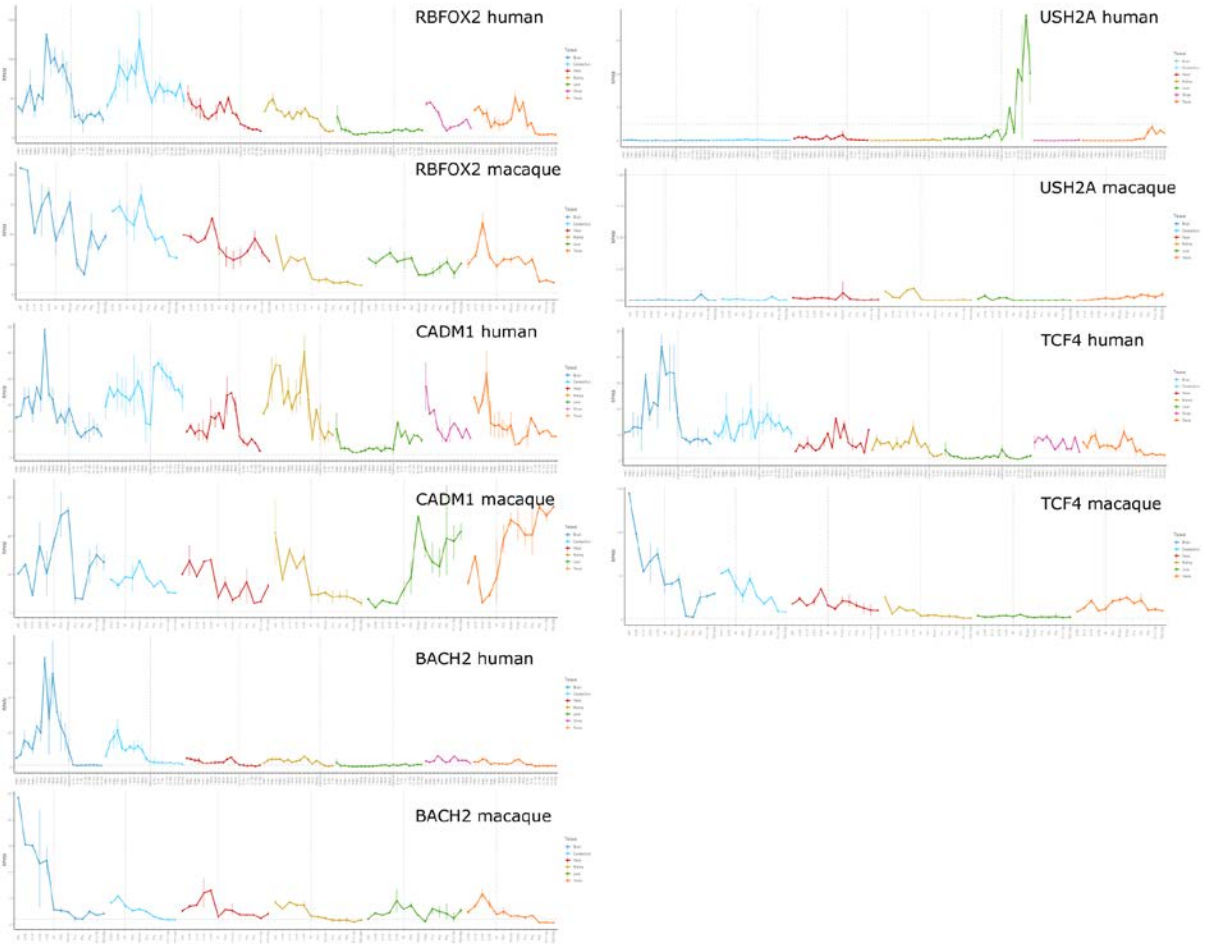


**Fig S13. Gene expression profiles across human and macaque organ development for the six genes overlapping HARs and extended gaps, with a recent modern human TMRCA (< 650 kya) and a deep modern human-Neanderthal TMRCA (≥ 650 kya; see Methods).**

**Table S1. *fastsimcoal* simulations of modern genomes with post-Out-of-Africa growth.** The table includes the ML parameter estimates, description of the template, and the estimate input file and block-bootstrap parameter estimates.

**Table S2. *fastsimcoal* simulations of modern genomes without post-Out-of-Africa.** The table includes the ML parameter estimates and a description of the template.

**Table S3. *fastsimcoal* simulations of the archaic genomes.** The table includes the ML parameter estimates, description of the template, and the estimate input file and block-bootstrap parameter estimates.

**Table S4. *fastsimcoal* simulations of modern genomes with post-Out-of-Africa growth, using the TSI genomes.** The table includes the ML parameter estimates and a description of the template.

**Table S5. *fastsimcoal* simulations of the introgressed Neanderthal.** The table includes the ML parameter estimates, a description of the template, and the estimate input file and block-bootstrap parameter estimates.

**Table S6. List of the 2,076 variants in Hardy-Weinberg Equilibrium deviation for excess of Neanderthal introgressed homozygotes, carrying at least one Neanderthal-specific derived allele.** Hardy-Weinberg exact test P, introgression rate, allele frequencies in the UKBBEUR, the Yoruba and the Altai genome, CADD PHRED score and correspondence with adaptive introgression signal found in Racimo et al. 2017 are reported.

**Table S7. NHGRI-EBI GWAS catalogue match with the clumped variants in HWE deviation for excess of Neanderthal homozygotes.**

**Table S8. GTEx match enrichment with the clumped variants in HWE deviation for excess of Neanderthal homozygotes compared to a DAF- and LDscore-matched genomic background.**

**Table S9. Constraints score comparison between expanded gaps, expanded introgressed regions and ILS.** Mean, standard deviation (SD), and pairwise comparisons using Wilcoxon signed-rank tests on the gene score per 10-kb window, controlled for the genic proportion of the window (see Methods).

**Table S10. Top 10-Mb merged windows for desertness.** We report the merged coordinates of 10-Mb windows in the top 1% and 5% for desertness, the average desertness computed on the top 1% and 5% windows, and their overlap with previously reported long desert.

**Table S11. Top 0.1% longest continuous expanded gaps.** The long gaps are annotated with average introgression rate, average recombination rate, number of overlapping Neanderthal-specific SNPs not observed in Yoruba genomes, region length, overlapping genes with the six genes involved in the significant filopodium-associated GO term (Fig. S11) in bold, and overlapping HARs.

**Table S12. HARs overlapping with expanded gaps evidencing a recent human TMRCA (<650 kya) and a deep modern human-Neanderthal TMRCA (≥ 650 kya).** The expanded gaps are annotated with the modern human, modern human-Neanderthal and modern-human-Denisova TMRCA, and HARs annotation directly taken from the original publication of the HARs used in this study, from Hubisz and Pollard, 2014 (see Methods). The position coordinates correspond to the 10-kb gap coordinates.

**Table S13. GTEx match enrichment with extended gaps associated with a recent modern human TMRCA (< 650 kya) and a deep modern human-Neanderthal TMRCA (≥ 650 kya).**
